# What does your gut tell you? Influences of a zombie-making and generalist fungal entomopathogen on carpenter ant micro- and mycobiota

**DOI:** 10.1101/2024.05.02.592155

**Authors:** Sophia Vermeulen, Anna M Forsman, Charissa de Bekker

## Abstract

Microbiome composition impacts many host aspects including health, nutrition, reproduction, and behavior. This warrants the recent uptick in insect microbiota research across species and ecosystems. Commensurate with this, the bacterial microbiome of the ant *Camponotus floridanus* has been well characterized across body regions and maturation levels. However, potential effects of entomopathogens on the gut microbiome, and the fungal communities therein, are yet to be assessed. Investigation of the microbiome during infection could provide insight into entomopathogenic infection and manipulation strategies and inform effective biopesticide strategies. Additionally, the mycobiome remains often overlooked despite playing a vital role in gut ecology with potential implications for health and infection outcomes. To improve our limited understanding of fungal infections in insects, and ants in particular, we characterized the effects of two entomopathogens with different infection strategies on the gut micro- and mycobiota of *C. floridanus* over time; *Ophiocordyceps camponoti-floridani* and *Beauveria bassiana.* Specialist, ‘zombie-making’ *O. camponoti-floridani* fungi hijack the behavior of *C. floridanus* ants over three weeks, causing them to find an elevated position, and fix themselves in place with their mandibles. This summiting behavior is adaptive to *Ophiocordyceps* as the ant transports the fungus to conditions that favor fruiting body development, spore production, dispersal, and transmission. In contrast, the generalist entomopathogen *B. bassiana* infects and kills the ant within a few days, without the induction of obvious fungus-adaptive behaviors. By comparing healthy ants with *Beauveria-* and *Ophiocordyceps-*infected ants we aimed to 1) describe the dynamics of the micro- and mycobiome of *C. floridanus* during infection, and 2) determine if the effects on gut microbiota are distinctive between fungi that have different infection strategies. While *Beauveria* did not measurably affect the ant host micro-and mycobiome, *Ophiocordyceps* did, especially for the mycobiome. Moreover, ants that were sampled during *Ophiocordyceps*-adaptive summiting behavior had a significantly different micro- and mycobiome composition compared to healthy controls and those sampled before and after manipulation took place. This suggests that the host microbiome might have a role to play in the manipulation strategy of *Ophiocordyceps*.

**Highlights:**

- *Ophiocordyceps* and *Beauveria* infections impact the gut microbiota of carpenter ants differently
- The fungal ASVs present in the ant gut are more diverse and dynamic across infection conditions compared to the bacterial ASVs
- There is significant dysbiosis in the bacterial and fungal microbiota of ants that display *Ophiocordyceps*-manipulated summiting behavior
- Dysbiosis in *Ophiocordyceps*-infected ants might be contributing to observed host behavioral changes while a comparable dysbiosis is not observed in *Beauveria* infections

## 1. Introduction

Parasitism has evolved multiple times across the fungal Kingdom, giving rise to numerous species that established antagonistic interactions with plants, animals, and other fungi. The order Hypocreales contains a multitude of parasitic species including many insect-infecting fungi (i.e., fungal entomopathogens) (1). The genera *Beauveria* (family: Cordycipitaceae) and *Ophiocordyceps* (family: Ophiocordycipitaceae) represent well-studied examples with different pathogenic strategies. *Beauveria bassiana* is a generalist that can infect insects across orders. As such, strains of this species are isolated and studied for their potential use in the biocontrol of insect pests (2, 3). To initiate infection, *Beauveria* spores attach and penetrate through the insect host’s cuticle. Once inside, the fungus proliferates as blastospores (i.e., single-celled, yeast-like cells), releases toxins, destroys host tissues, and kills the insect within a few days (3). This destructive, toxin-based infection strategy, paired with its wide host range classify *Beauveria* as a necrotroph-like fungal entomopathogen.

In contrast to *Beauveria*, *Ophiocordyceps* species are highly specialized entomopathogens that generally infect only one insect species (4–6). The ‘zombie-making’ fungi of the species complex *Ophiocordyceps unilateralis* sensu lato infect ants from the Camponotini tribe. These fungi slowly hijack ant behavior over the course of three to four weeks before eventually killing their host (7, 8). While the fungus is growing as single blastospores, infected individuals wander away from the nest and climb to elevated positions that provide favorable conditions for fungal growth (9, 10). The manipulated ant then proceeds to bite onto the vegetation, firmly attaching itself in the final moments before death. Following this summiting behavior, the fungus converts to a multicellular, mycelial growth, which eventually gives rise to a fruiting body that sprouts from the cadaver (11, 12). This fruiting body carries the spores that are released to infect new ants (4–6). These manipulated behaviors are a crucial aspect of the *Ophiocordyceps* life cycle that aid in circumventing social immunity behaviors of healthy nest mates (13) and place the ant in a microclimate that promotes fungal growth and spore dispersal (10, 14). Without this behavior, the fruiting body cannot form (11, 15), and transmission fails. Therefore, *Ophiocordyceps* species within the *unilateralis* complex must treat their host in a much more delicate manner than *B. bassiana* to ensure that the ant does not succumb to the infection prematurely. As such, *Ophiocordyceps* is thought to begin its infection much like a biotrophic fungal plant pathogen, gaining resources from living cells, and only inducing subtle changes without causing severe tissue damage. However, once manipulated biting occurs, the fungus necessarily becomes more necrotrophic as it rapidly needs to consume insect tissue to gain enough energy for fruiting body development. As such, in contrast to *B. bassiana*, ant-infecting *Ophiocordyceps* fungi use parasitic strategies comparable to those of a hemi-biotroph (16).

Unraveling how *Ophiocordyceps* and other insect-infecting fungi interact with their insect hosts could prove valuable for the discovery of novel, more sustainable pesticides, and pesticide targets. Moreover, learning how pathogenic behavioral phenotypes are induced leads to a better mechanistic understanding of insect behavior in general. As such, recent laboratory studies on *Ophiocordyceps*-ant interactions have used a combination of quantitative behavioral assays and various omics technologies to investigate how *Ophiocordyceps* affects ant circadian activity, communication, olfaction, immunity, and biogenic amine levels such as dopamine (8, 13, 17–19). The fungus appears to achieve this by secreting small bioactive molecules of which some are predicted to bind to G-protein coupled receptors that are involved in light and odor sensing or binding of biogenic amines (20). While these mechanistic studies are vital to understand how *Ophiocordyceps* manipulates its host through chemical means, they only focus on fungal interactions with the nervous and muscular tissue in the ant head. This approach leaves the interactions between the fungus and other host tissues, such as the gut and its microbiota, completely in the dark. Meanwhile, microbiome studies produce mounting evidence for the importance of (gut) microbiota in general host functioning, including nutrition, immunity, and behavior (21–25). As such, interactions between infectious agents, such as fungal entomopathogens and other parasites, and the gut microbiota of their host could be a vital part of disease progression and resulting phenotypes (22, 26). This could be especially true for fungal specialists like *Ophiocordyceps* that rely on altered behavioral phenotypes to increase their fitness outcomes. Most studies that examined the microbiota–brain axis so far have focused on mouse models to demonstrate links between gut dysbiosis and neurodegenerative diseases, emotional state and social disfunctions (27–31). Nevertheless, *Drosophila* models have also been able to link gut dysbiosis with changes in short-chain fatty acids, which are considered an Alzheimer’s marker (32, 33). In addition, studies with honeybees produced results that corroborate findings obtained in autism-spectrum disorder research with mice and humans (24). Furthermore, the gut microbiome of *Drosophila* has been linked to octopamine-dependent aggression behaviors (34), while ant gut bacteria have been suggested to impact chemical communication and, subsequently, olfactory-dependent social behaviors (35).

While there is previous work that connects insect pathologies and behavior with microbiota composition, the majority of microbiome research, including studies done on ants, focuses on comparisons between species, ecosystems, diet composition, and social status (25, 36–42). Most of these studies have only focused on the bacterial microbiota, leaving the fungal microbiome (i.e., mycobiome) underexplored. Studies that have investigated insect mycobiomes suggest that fungal communities can also be impacted by disease and show different patterns in community composition when compared to the bacterial microbiome (43). Both the bacterial and fungal microbiome of grain beetles infected with tapeworms showed changes when compared to uninfected individuals (44). Moreover, the gut mycobiome of honeybees was shown to be much more variable than their bacterial microbiome (36) and together with the bacterial gut community to be responsible for shaping social status (39). In fact, mycobiome explorations in mosquitoes demonstrated that interkingdom interactions between fungal and bacterial microbiota shape these communities, with consequences for host growth and development (45). These studies suggest that research into insect mycobiomes would provide a more complete picture of the microbiome while potentially revealing a different side of microbiome-host interactions. Moreover, studies that investigate the effects of infectious diseases on the gut microbiota of vertebrates suggest their involvement in disease outcomes (46–48). With regards to the topic of fungal pathogens of insects, the intersection between insect pathology and the microbiome is even considered to be an avenue of research that may improve the use of these fungi in the biocontrol of insect pests (26).

Currently, the interaction between the behavior-manipulating fungus *Ophiocordyceps camponoti-floridani* and its carpenter ant host *Camponotus floridanus* is one of the more in-depth studied examples of summit disease (8, 10, 13, 19, 20, 49). Additionally, the bacterial microbiome of *C. floridanus* has been well characterized across different body regions (25) and maturation levels (50). Nevertheless, the fungal communities and effects from disease on *C. floridanus* microbiota have yet to be assessed. Since microbiome composition has been linked to behavior in other insects, and infection can seemingly change that composition, we propose that part of the behavioral changes observed in *Ophiocordyceps-*manipulated ants may be attributed to their microbiota. As such, we set out to investigate the bacterial and fungal gut microbiomes of *Ophiocordyceps*-infected *C. floridanus* using DNA metabarcoding. We compared our findings to data obtained from *Beauveria*-infected ants to answer if 1) fungal infection alters the bacterial and fungal community composition of the gut microbiome in *C. floridanus* when compared to healthy nestmates, 2) the gut microbiome community changes as disease progresses, and 3) the microbiome community is affected differently depending on the infecting entomopathogen (*Ophiocordyceps* vs *Beauveria*). We hypothesized that the microbiome of infected ants would diverge from that of healthy nestmates, and that *Ophiocordyceps* and *Beauveria* infections would affect microbiome compositions differently since they are fungi with contrasting parasitic lifestyles. If supported, this would suggest that changes in the host gut microbiota caused by *Ophiocordyceps* infection could underly some of the behavioral changes observed.

## 2. Methods

### 2.1 Ant collection and housing

To conduct our metabarcoding analysis of carpenter ant guts upon fungal infection, we collected two *C. floridanus* colonies from the University of Central Florida’s (UCF) arboretum (Colony 1 and 2) and two additional colonies from the Chuluota Wilderness area (Colony 3 and 4). We received permits for these collections from the UCF Arboretum and Seminole County Natural Lands, respectively. All collected colonies were queenless and averaged between one to two thousand individuals consisting of majors, minors, and brood.

Upon collection, we placed each colony into a 9.4 L plastic container (Rubbermaid) with *ad libitum* water and food (i.e., frozen crickets and 15% sugar water). Several darkened 50 mL tubes (Greiner) served as nest chambers and talcum-lined walls prevented the ants from climbing out. To reset *C. floridanus* biological clocks to lab conditions, we placed the colonies in an incubator (I36VL, Percival) under constant light, 25 °C temperature, and 70% relative humidity for 48 hours. Subsequently, we entrained the colonies to 24h rhythms of light and temperature (i.e., environmental entrainment cues known as Zeitgebers). We used a 12 hr :12 hr light-dark cycle at 28°C and 20°C, respectively, and 70% relative humidity. The cycle began with lights on at Zeitgeber time (ZT) 0 and followed a four-hour increasing ramping step to full light and 28°C at ZT 4. These settings were followed by a four-hour hold at peak light and temperature until ZT 8, and a four-hour decreasing ramp step to complete darkness and 20°C at ZT 12.

Seven days prior to infections, we separated two groups of 110 minor caste workers from each colony into new 9.4 L plastic containers with a plaster bottom. *Tillandsia* is the main biting substrate for *Ophiocordyceps*-infected *C. floridanus* in Central Florida (Will et al., 2023). To provide this substrate for manipulated individuals to latch onto, we draped *Tillandsia* on two upright 12 cm wooden skewers embedded in the plaster on one end of the container. On the other end of the container, we placed a solid dark plastic bottom of a micropipette tip box (TipOne) upside down to serve as a nest area. We fed all fragment colonies ad libitum on 15% sucrose solution and water throughout the experiment.

### 2.2 Fungal culturing and ant infections

To infect carpenter ants with a behavior-manipulating hemi-biotroph-like fungus, we used *O. camponoti-floridani* strain OJ 2.1, isolated in January 2021 from a naturally infected *C. floridanus* ant cadaver found in Ocala National Forest (Florida). Seven days prior to infections, we started an 8 mL liquid culture by inoculating Grace’s Insect Medium (Invitrogen) supplemented with 2.5% Fetal Bovine Serum (FBS) (Invitrogen) with blastospores obtained from a frozen glycerol stock. We incubated the culture at 28 °C in the dark at 50 rpm. For necrotroph-like fungal infections we used *B. bassiana* strain ARSEF 2860. To obtain fresh conidiospores, we inoculated potato dextrose agar (PDA) by rolling cryobeads containing frozen spores across the surface. For blastospore growth, we harvested the fresh spores from a fully colonized PDA plate and inoculated a 20 mL liquid culture of Sabouraud dextrose broth (Sigma) six days prior to infections. We incubated liquid cultures at 25 °C in the dark at 120 rpm.

To conduct *Ophiocordyceps* and *Beauveria* infections, we harvested fungal blastospores immediately prior to injections by filtering the cultures through sterile gauze. Following centrifugation and supernatant removal, we resuspended the cells in fresh Grace’s media supplemented with 2.5% FBS. *Ophiocordyceps* was resuspended to a cell concentration of 4.6 x 10^7^ blastospores/mL, while the *Beauveria* infection solution was brought to 3.5 x 10^7^ blastospores/mL. We performed infections with 10 µL borosilicate capillary tubes (Drummond) pulled to a needle point with a PC-100 Narishige instrument.

To establish *Ophiocordyceps* infections, we injected two times 110 ants, obtained from two different colonies (colony 1 and 2), on the ventral side of the thorax under the first pair of legs with 0.5 uL of blastospore solution. The healthy control groups also consisted of two times 110 ants, taken from the same two colonies, which were sham-injected with 0.5 mL Grace’s media supplemented with 2.5% FBS without fungal cells (Will et al., 2020). Following injection, we left ants undisturbed for three days after which we removed deceased ants, considering them a result of injection trauma. As a result, 79% of *Ophiocordyceps*-injected and 85% of sham-injected ants survived, which were kept in the infection experiment for further observation, data collection and sampling.

While injection of fungal blastopores is necessary for *Ophiocordyceps* infection, *Beauveria* infections are best established by pricking ants with a capillary needle that has been dipped briefly into a fungal blastospore solution (13). As such, we conducted *Beauveria* infections in two times 110 ants, again obtained from two different colonies (colony 3 and 4), by pricking them on the ventral side of the thorax under the first pair of legs with a blastospore-coated capillary needle. In this case, the control groups of two times 110 ants, taken from the same colonies, were pricked with a needle dipped in a sham solution of Grace’s-FBS 2.5% without fungal cells. We left ants undisturbed for one day after which we removed deceased ants and counted all surviving ants as part of the experiment, which comprised 97% of *Beauveria*-pricked and 98% of sham-pricked ants.

### 2.3 Daily observations, survival data collection and ant sampling

After leaving *Ophiocordyceps-*infected ants undisturbed for three days post injection (DPI), we checked survival and removed dead individuals daily between ZT 5 and 7 (i.e., while lights were on). At 19 DPI, 25% of the infected population had died, which we indicated as lethal time (LT) 25 (Figure 1A). At this time, none of the infected individuals had shown manipulated summiting and biting behaviors yet. However, most individuals that make it across this time point eventually progress towards the ultimate summiting behavior (8). As such, to investigate potential microbiome changes prior to manipulated biting, we included LT25 as our first sampling time point (Figure 1A).

**Figure 1:**
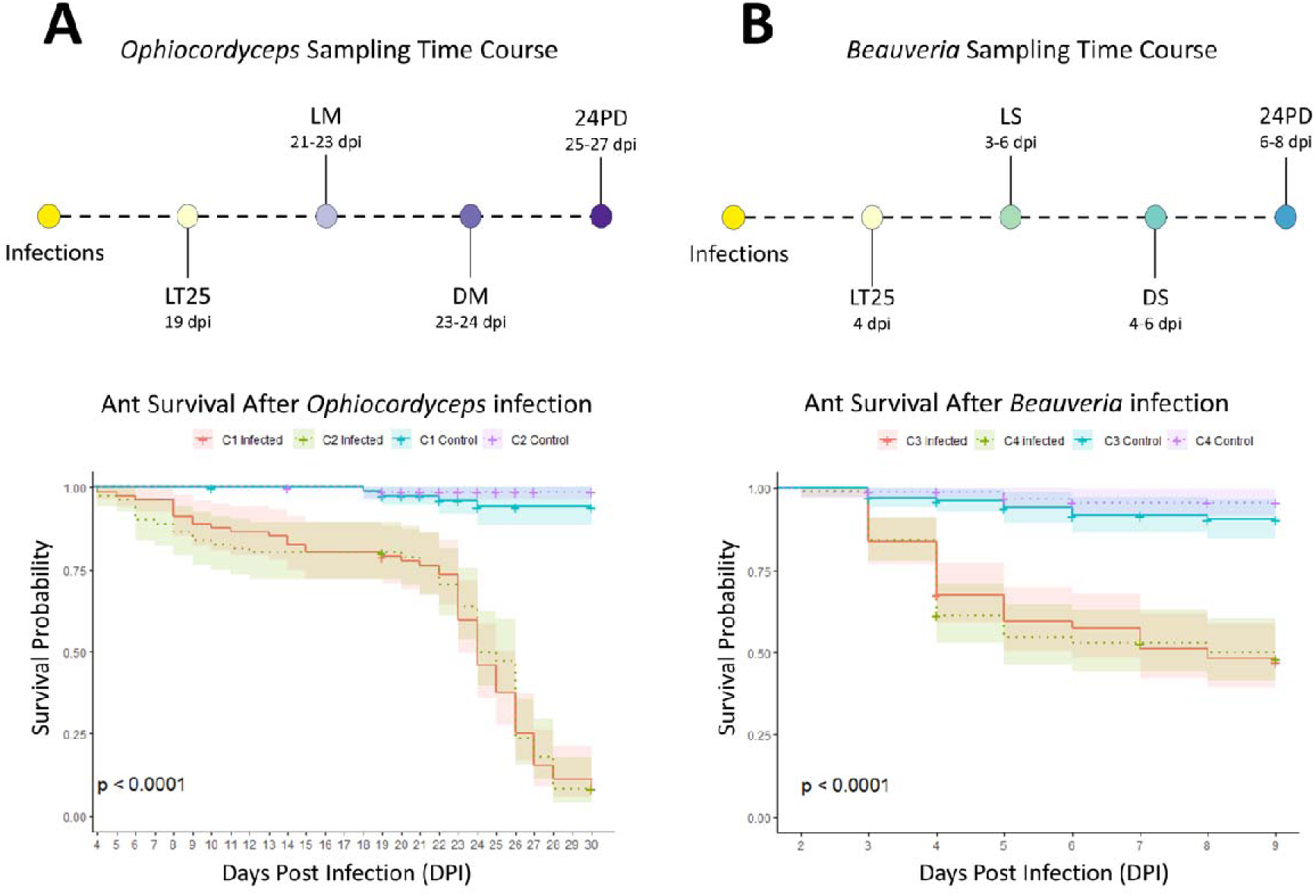
Sampling regime and survival curves of C. floridanus ants infected with O. camponoti-floridani or B. bassiana. Visualization of the parallel sampling timelines that were used to collect A) Ophiocordyceps-infected and B) Beauveria-infected ants for this study. The corresponding survival curves for both these infections are plotted underneath these timelines, demonstrating that both infections led to a significant reduction in survival probability for both infections as compared to healthy control ants obtained from the same colony (p < 0.0001). For both infections two independently collected ant colonies were used: colony 1 and 2 (C1 and C2) were used for Ophiocordyceps infections and colony 3 and 4 (C3 and C4) were used for infections with Beauveria. The two colony replicates for each infection showed a comparable survival probability. Samples were collected at the time when 25% of the ants had died (i.e., lethal time LT25), ants showed manipulated or sickness behavior (i.e., live manipulated LM and live sick LS), ants had died after showing these behaviors (i.e., dead manipulated DM and dead sick DS), and 24 hours after the ant had succumbed from the respective infections (i.e., 24 hours post death 24PD).

For our second *Ophiocordyceps* infection time point, we wanted to include live individuals that displayed the characteristic manipulated summiting and biting behavior (i.e., “Live Manipulation” or LM). Previous studies on *O. camponoti-floridani* infections found that manipulated biting occurs around ZT 21 under laboratory conditions (Will et al., 2020). So, as we neared the three-week mark when these manipulations typically begin, we also made daily observations around ZT 21-23 to catch ants in the act of summiting and biting. At 21 DPI, the first manipulations occurred (Figure 1A). We observed ants overnight, checking every two hours and found manipulations started between ZT 18 and 22, but could be as late as ZT 1 in one case. We did not collect ants until ZT 22 to avoid disturbing other ants that might be climbing and biting. At this time, live manipulated ants were still very responsive (i.e., moving legs and antennae rapidly when gently poked with forceps).

After manipulation, *Ophiocordyceps* is thought to rapidly switch its biotroph-like growth towards a more necrotrophic one as it begins to switch from single-celled blastospores to a multicellular hyphal growth, killing the ant host in the process (7). To capture potential microbiome changes during this process, for our third time point, we included ants that had just succumbed to the *Ophiocordyceps* infection after displaying manipulated summiting behavior (i.e., “Dead Manipulation” or DM) (Figure 1A). Ants for this time point were collected by conducting hourly checks following manipulation to monitor time of death, which occurs within a few hours upon manipulated biting. We considered ants dead when they no longer moved in response to being poked with forceps.

For our fourth timepoint, we included ants 24h after manipulation and death when tissue damage due to cell death and fungal carbon and nutrient consumption is more abundant (i.e., “24Hr Post Death” or 24PD (Figure 1A). To collect these samples, we moved dead manipulated individuals to an isolated container with an identical setup to the original one, kept within the same incubator for 24h. Isolation protected the dead ants from being removed/dismembered by their live nestmates as part of their social immune response.

To collect *Beauveria-*infected ants we aimed to collect comparable timepoints to the *Ophiocordyceps* experiment. However, *Beauveria’s* much more rapid disease progression required a different monitoring and sampling approach. After leaving ants undisturbed for one day, we conducted survival and behavioral observations every 4 hours. As we neared the LT25 timepoint, observations were done every two hours to ensure precise timing. Because ant mortality at this point was a result from infection followed by rapid death, there was necessary overlap in the LT25 sample collection and later timepoints (i.e., ‘sick’ and “dead”). The *Beauveria* infection experiment reached LT25 at 4 DPI (Figure 1B). To distinguish this first sampling group (LT25) from the second collection timepoint (the ‘sick’ phenotype), only ants that were out of the nest and able to walk at LT25 were selected.

Since *Beauveria* does not manipulate host behavior, we collected *Beauveria-*infected ants at a distinct “sick” phenotype, starting at 3 DPI (i.e., “live sickness behavior” or LS) (Figure 1B). Ants that displayed the “sick” phenotype could no longer walk more than a few steps without falling but still actively responded to a poke with forceps. They were often found on their sides while moving their legs but unable to stand. We considered this a comparable timepoint to the *Ophiocordyceps* ants collected at live manipulation because once *Beauveria-*infected ants displayed this behavior, they similarly died within a few hours (Figure 1B).

To collect *Beauveria*-infected “dead” ants we isolated “sick” ants in identical container setups to prevent disturbance from nest mates and monitored them hourly until death, after which they were immediately sampled (i.e., “dead sickness behavior”, or DS). To collect *Beauveria*-infected ants 24h post death (24PD), we left expired ants that had displayed the “sick” phenotype in isolation for 24h under the same incubator conditions as the remainder of the experiment (Figure 1B).

We sampled ten *Ophiocordyceps*-infected ants, ten *Beauveria*-infected ants, and ten sham-injected and -pricked control ants at each timepoint (i.e., LT25, LM/LS, DM/DS, and 24PD). Three additional *Ophiocordyceps*-infected ants were collected at live manipulation due to availability. Eventually, we reduced the control group to comprise twelve healthy sham samples randomly selected from across the four *Ophiocordyceps-*infection timepoints and twelve from the *Beauveria*-infection timepoints. To maintain microbiome integrity, we flash froze ants in liquid nitrogen immediately following collection.

### 2.4 DNA extraction, library preparation, and sequencing

To conduct bacterial 16S and fungal ITS metabarcoding on healthy and infected ant guts, we processed all samples in a randomized order. First, we separated gasters from frozen ants under sterile conditions and placed them in 2 mL Eppendorf tubes with two metal ball bearings (5/32” type 2B, grade 300, Wheels Manufacturing). We disrupted the cells in frozen gasters by placing the tubes in a Mini G Tissue Homogenizer (SPEX) for 60 seconds at max rpm. We extracted DNA from these samples using the ZymoBIOMICS DNA microprep kit (Zymo Research, D4301) following manufacturer’s recommendations. Using the extracted DNA, we prepared amplicon libraries according to the 16S Metagenomic Sequencing Library Preparation protocol from Illumina. We also included negative DNA extraction controls to account for potential contaminants that we introduced into our samples during this process.

For bacterial metabarcoding, we targeted the V4 region of the 16S SSU rRNA gene using primers 515F (51) and 806R (52). The internal transcribed spacer (ITS) region is predominantly used for fungal metabarcoding (53, 54). As such, we targeted the ITS2 region using ITS primers ITS3_KYO2 (54) and ITS4 (55). We conducted amplicon PCR (PCR1) in 25 µL reactions using 2.5 µL of DNA sample, 5 µL each of 1 µM forward and reverse primer, and 12.5 µL of KAPA HiFi HotStart ReadyMix (Roche). We ran the 16S amplicon reactions in duplicate using the following PCR conditions: initial denaturation at 95°C for 3 minutes, followed by 25 cycles of 95°C for 30 seconds, 50°C for 30 seconds and 72°C for 30 seconds, and a final extension at 72°C for 5 minutes. We performed the ITS reactions in singleton using the same PCR conditions, with the exception that the annealing temperature used was 47°C. Following PCR, we cleaned PCR1 products by removing primer dimers with 1.5x Sera-Mag SpeedBead solution following the protocol described in (56). The duplicate 16S PCR1 reactions were combined during this process. We also included negative PCR1 controls to account for potential contaminants we introduced into our samples.

Prior to the index PCR (PCR2), we checked the fragment size of randomly selected 16S and ITS libraries using a Tapestation 4200 (Agilent) and D1000 HS reagents (Agilent). For both 16S and ITS, we detected expected fragment sizes between 300-400 bp. We uniquely dual-indexed all ITS libraries in 50 µL PCR reactions, consisting of 8 µL PCR1 product, 5 µL each of Nextera XT Index primers i5 and i7 (Illumina), 7 µL molecular grade water (Invitrogen), and 25 µL KAPA HiFi HotStart ReadyMix (Roche). The 16S libraries were indexed in 25 µL PCR reactions consisting of 8.5 µL PCR1 product, 2 µL each of Nextera XT Index primers i5 and i7, and 12.5 µL KAPA HiFi HotStart ReadyMix. We used the following PCR conditions for both the ITS and 16S libraries: initial denaturation at 95°C for 3 minutes, followed by 8 cycles of 95°C for 30 seconds, 55°C for 30 seconds and 72°C for 30 seconds, and a final extension at 72°C for 5 minutes. We cleaned PCR2 products with magnetic beads at a 1.2x concentration to remove smaller base pair fragments following the protocol as described for PCR1.

Following PCR2, we quantified all libraries using a Qubit fluorometer with the 1X dsDNA HS assay kit (Invitrogen) to create an equimolar pool for sequencing. After an additional 1.2x magnetic bead cleanup, we created a high and a low concentration pool, quantified the two pools using an NEBNext Library Quant Kit for Illumina (New England Biolabs), and determined average fragment size and adaptor dimer removal on a Tapestation 4200 (Agilent) with D1000 HS reagents (Agilent). Subsequently, we created a final pool by combining the 16S and ITS library pools in equimolar amounts for sequencing using an Illumina MiSeq instrument, sourced by the UCF Genomics and Bioinformatics Cluster, with a paired-end approach (2×300bp) and V3 reagents (Illumina).

### 2.5 Sequence data processing

We demultiplexed sample reads on the MiSeq instrument prior to importing paired-end read data into QIIME2 (57) for further processing and quality control. Using the QIIME2 *cutadapt trim-paired* function, we removed reads lacking primer regions and trimmed forward and reverse primer regions from reads that did have them. We then proceeded to denoise, dereplicate, and merge the trimmed forward and reverse reads using the *dada2 denoise-paired* function to create amplicon sequence variants (ASVs).

For ITS reads, we performed denoising with and without truncation (QC or otherwise) to determine the most optimal procedure for our data because the fungal ITS region can be rather variable in length. As such, it is generally not recommended to trim reads at the 3’ end to prevent bias in amplicon size (58). However, not trimming reverse reads left many taxa unidentified after the taxonomy assignment. Similarly, we attempted to process ITS forward reads following the same procedures for single-end data without truncation. This still left many taxa unidentified. The ITS reads with truncation *(trunc-len-f 276; trunc-len-r 173)* proved to have the most classified reads compared to the other methods. Therefore, we used truncated data for downstream analyses while acknowledging that doing so may have introduced some bias in amplicon size.

We taxonomically classified 16S ASV’s using the Green Genes 13.5 database at 99% identity with BLAST+ (QIIME2, *classify-consensus-blast*; (59) and created a phylogenetic reference tree with a Green Genes 13.8 SATé-enabled phylogenetic placement database (QIIME2, *fragment-insertion sepp*; (60). We classified ITS ASV’s using a naïve bayes classifier (QIIME2, *fit-classifier-naive-bayes*) trained on the fungal 9.0 UNITE database (61). After assigning taxonomy, we filtered out those 16S reads not identified to at least the phylum level, those assigned as being of chloroplast or mitochondrial origin, and those that did not meet the set minimum of three reads per ASV. Rarefaction curves (package: vegan – *rarecurve*) indicated that a sequencing depth of 4000 was sufficient to capture most ASVs. Of all our collected samples, 94 passed this cutoff (extraction and PCR controls not included) (Supplementary File 1, Figure 1 and Table 1). We filtered ITS reads similarly, excluding reads that were not identified to the phylum-level and requiring a minimum of three reads per ASV. We also filtered out reads identified as *Ophiocordyceps spp.* and *Beauveria spp.* as these were the infecting entomopathogens. Rarefaction curves indicated sufficient ASV coverage at a sequencing depth of 500. Out of all our samples, 81 samples passed this cutoff (extraction and PCR controls not included) (Supplementary File 1, Figure 1 and Table 1).

### 2.6 Statistical analyses

All statistical analyses were conducted in R version 4.2.1. We analyzed survival probability of ants across the infection period with the R package survival version 3.5-8 (62) and constructed the survival curves using survminer version 0.4.9 (63).

To determine alpha and beta diversity of overall biological (infection) groups and separate infection timepoints, we paired classified reads and taxonomy data with sample metadata and imported this into R as phyloseq objects. We calculated within-sample alpha diversity using observed richness and Shannon diversity (package: phyloseq – *estimate_richness*) (64). To test for differences between diversities, we used a Kruskal-Wallis rank sum test, followed by pairwise comparisons using the Dunn test (package: dunn.test - *dunn.test*, p-value adjustment method for multiple comparisons = Bonferroni) (65). We visualized alpha diversity differences by generating box plots using ggplot2 (66).

To detect differences between groups and timepoints we also used beta diversity metrics and performed pairwise permutational multivariate analyses of variance (PERMANOVA) (package: pairwiseAdonis – *pairwise.adonis,* method = bray curtis, permutations = 999, p-value adjustment method for multiple comparisons = Bonferroni) (67). We visualized beta diversity using non-metric multidimensional scaling (NMDS) ordination of Bray-Curtis distances (package: phyloseq – *plot_ordination*) (64). We produced taxa bar plots using ggplot2 (66).

Using these methods, we compared healthy control ants that were obtained from the same colonies as the *Ophiocordyceps* infection and the *Beauveria* infection and found that their microbiota had no significant differences in alpha or beta diversity. Therefore, we combined them into a single healthy control group for comparisons to the infected ant groups. Moreover, this indicated that the ants used in these infection experiments had similar microbiota despite being from different colonies and field collections, allowing for direct comparisons between the two fungal infection treatments.

To determine whether the type of fungal infection (i.e., necrotroph-like versus hemi-biotroph-like) affected the bacterial and fungal communities of the gut microbiome differently, we grouped reads from timepoints LT25 through 24HR into two categories based on the infecting entomopathogen (treatment groups): *Ophiocordyceps-* or *Beauveria-*infected ants. For timepoint comparisons, we separated reads from the two fungal groups back into their respective timepoints: control, LT25, LM/LS, DM/DS, 24PD. In doing so, we discovered one outlier ITS *Beauveria* 24HR sample that had high NMDS1 and NMDS2 scores (Supplementary File 1, Figure 2). After the filtering steps, and removal of *Ophiocordyceps* and *Beauveria* reads, this sample was only left with unidentified reads and reads identified as *Malassezia*, a likely contaminant. Therefore, we removed this sample from both alpha and beta diversity analyses.

**Figure 2:**
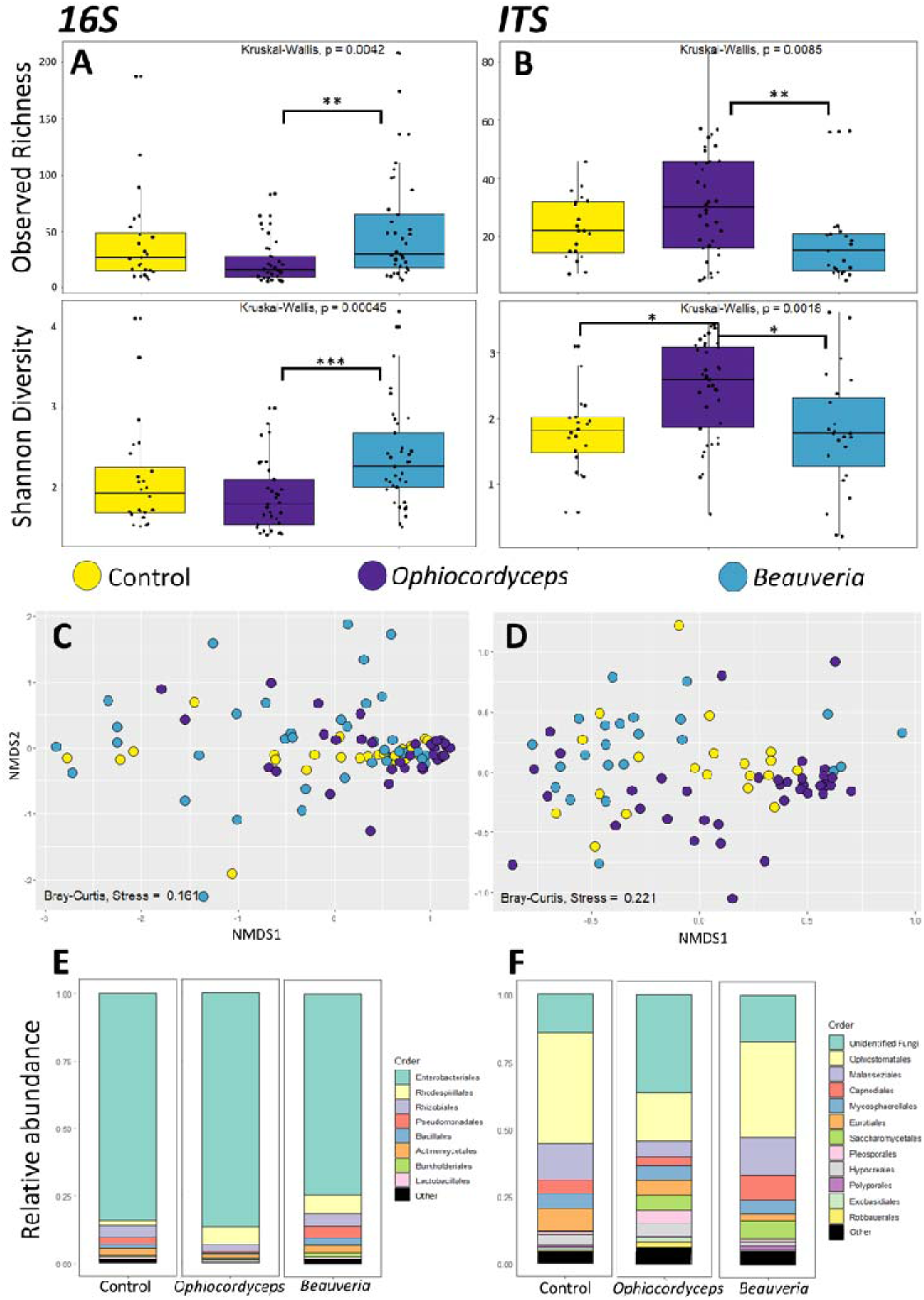
Differences in bacterial microbiota and fungal mycobiota between healthy C. floridanus ants and those infected with O. camponoti-floridani or B. bassiana. Box plot representations of the bacterial (16S) (A) and fungal (ITS) (B) alpha diversity in the guts of healthy control ants, ants infected with Ophiocordyceps, and ants infected with Beauveria. Adjusted p-values of significantly different observed ASV richness and Shannon diversity between infection groups are indicated as follows: * p<0.05, ** p< 0.01, *** p<0.001. (C and D) Beta diversity is depicted in non-metric multidimensional scaling (NMDS) plots for the different infection treatment groups. The points represent bacterial (D) and fungal (E) communities from individual gut samples, and their color indicates which treatment group each sample belongs to. C) NMDS plot using a Bray Curtis distance matrix of the bacterial gut microbiome for different infection groups (stress = 0.161). D) NMDS plot using a Bray Curtis distance matrix of the fungal gut mycobiome for different infection groups (stress = 0.22). E and F) Taxa bar plots showing the relative abundance of bacterial (E) and fungal (F) orders present in the guts of healthy control ants, ants infected with Beauveria, and ants infected with Ophiocordyceps.

## 3. Results

### 3.1 Overall data quantity and quality

Sequencing of ant samples resulted in 6,670,956 paired-end reads across all samples for 16S libraries and 5,968,530 paired-end reads across all samples for ITS libraries. After filtering, taxonomic assignment, and rarefaction, we retained 1280 ASVs across 94 samples for 16S libraries and 833 ASVs across 81 samples for ITS libraries (Supplementary File 1, Table 1).

To monitor and account for contamination introduced during DNA extraction and library preparation, we included negative controls for DNA extraction (extraction controls) and library preparation (PCR controls). Both technical control types had a read coverage similar to biological samples and detected comparable numbers of taxa. Consequently, we could not reliably use these controls to remove potential contaminants from our samples. To investigate the source of contaminant introduction, we grouped samples by DNA extraction date and PCR 1 date and analyzed alpha diversity metrics (Observed richness, Shannon). While certain extraction dates were significantly different, there was no gradual increase in richness or Shannon diversity over time (Supplementary File 1, Figure 3). This indicates that the reagents themselves were not contaminated during the process and that the taxa observed in the negative controls were likely introduced due to some level of cross-contamination during extraction. Grouping samples by PCR1 date did not show any significant differences (Supplementary File 1, Figure 4), indicating that no measurable additional cross-contamination was introduced during library preparation. Taken together, our biological samples have some level of cross-contamination that might obscure present differences between groups. However, because we processed all samples in a random order (both extractions and PCR reactions), any significant differences detected between treatment groups or infection time points cannot be attributed to batch effects from the days they were processed.

**Figure 3:**
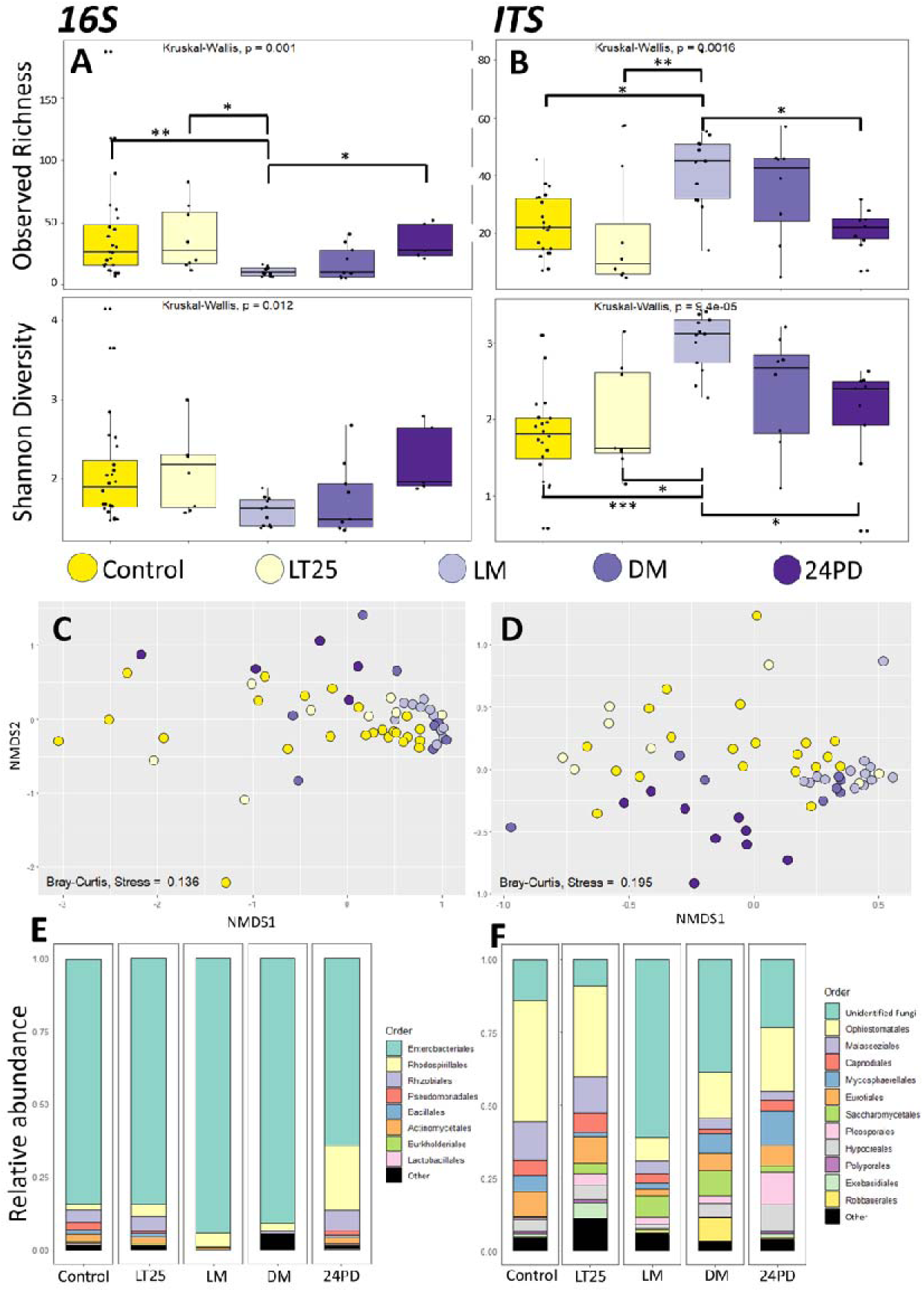
Differences in bacterial and fungal gut microbiota of C. floridanus during infection progression and manipulation by O. camponoti-floridani. Box plot representations of bacterial (16S) (A) and fungal (ITS) (B) alpha diversities in the guts of healthy control ants and ants infected with Ophiocordyceps, which were sampled before (LT25), during (LM) and after manipulated summiting and biting behavior (DM and 24PD). Adjusted p-values of significantly different observed ASV richness and Shannon diversity between infection groups are indicated as follows: * p<0.05, ** p< 0.01, *** p<0.001. C and D) Beta diversity is depicted in non-metric multidimensional scaling (NMDS) plots for the different Ophiocordyceps infection time points. The points represent bacterial (D) and fungal (E) communities from individual gut samples, and their color indicates which infection time point each sample belongs to. C) NMDS plot using a Bray Curtis distance matrix of the bacterial (16S) gut microbiome for different infection time points (stress = 0.135). D) NMDS plot using a Bray Curtis distance matrix of the fungal (ITS) gut mycobiome for different infection time points (stress = 0.195). E and F) Taxa bar plots at order level showing the relative abundance of bacterial (E) and fungal (F) microbiota present in the guts of healthy control ants, and ants infected with Ophiocordyceps prior (LT25), during (LM) and after manipulation (DM and 24PD).

**Figure 4:**
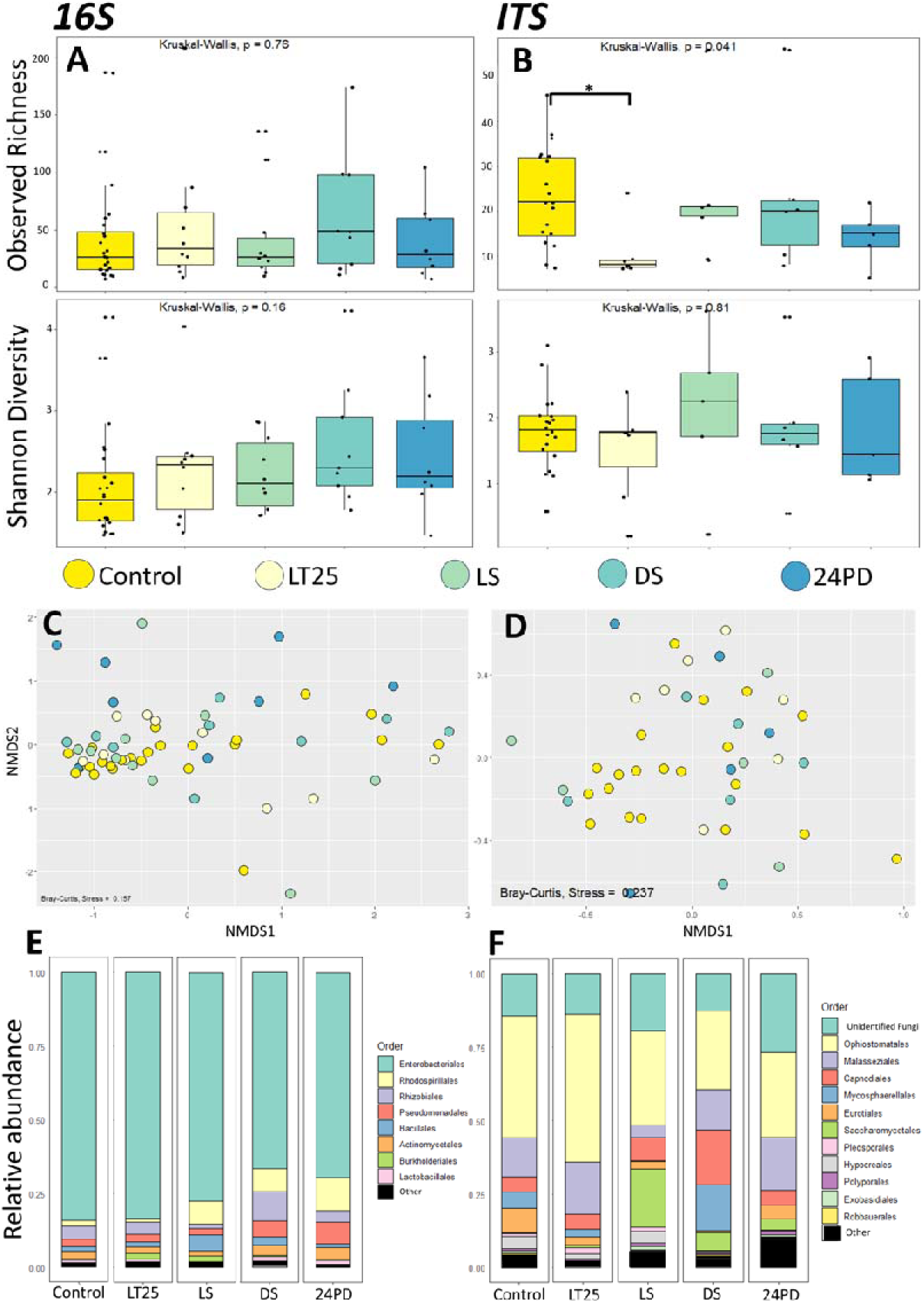
Bacterial and fungal gut microbiota of C. floridanus during infection by B. bassiana. A and B) Box plot representations of the bacterial (16S) (A) and fungal (ITS) (B) alpha diversity within the guts of healthy control ants and ants infected with Beauveria, which were sampled at different time points during infection. The adjusted p-value of the sole significantly different observed ASV richness between 24 hour after death (24PD) and healthy control is indicated as * p<0.05. C and D) Beta diversity is depicted in non-metric multidimensional scaling (NMDS) plots for the different Beauveria infection time points. The points represent bacterial (D) and fungal (E) communities from individual gut samples, and their color indicates which infection time point each sample belongs to. C) NMDS plot using a Bray-Curtis distance matrix of the bacterial (16S) gut microbiome for different infection time points (stress = 0.159). D) NMDS plot using a Bray-Curtis distance matrix of the fungal (ITS) gut mycobiome for different infection time points (stress = 0.236). E and F) Taxa bar plots at order level showing the relative abundances of bacterial (E) and fungal (F) taxa present in the guts of healthy control ants, and ants infected with Beauveria prior to displaying a “sick” phenotype (LT25), during (LS) and after displaying the “sick” phenotype (DS and 24PD).

### 3.2 Differences in alpha diversity between infection treatments

When we compared alpha diversity of gut microbiota in healthy ants to those infected with *Ophiocordyceps* and *Beauveria*, we did not find an effect of fungal infection on bacterial (16S) alpha diversity, irrespective of infection time point (Figure 2A). However, the mycobiome (ITS) in *Ophiocordyceps-*infected ants had a significantly higher Shannon diversity index (p = 0.01) than healthy control ants (Figure 2B, Supplementary File 2, Table 2). These results indicate that *Ophiocordyceps* infection may affect the relative abundance of fungal taxa present within the healthy ant host gut.

When we compared the effects of *Ophiocordyceps* and *Beauveria* infections, we found that the alpha diversities of both bacterial microbiome and fungal mycobiome differed significantly among infection groups (Figure 2A and B). The bacterial gut microbiome of *Ophiocordyceps*-infected ants had a significantly lower observed ASV richness (p=0.003) and Shannon diversity index (p = 0.0003) than those infected with *Beauveria* (Figure 2A, Supplementary File 2, Table 2). However, we observed the reverse for the fungal gut mycobiome; the mycobiome of ants infected with *Ophiocordyceps* had a significantly higher observed richness (p = 0.006) and Shannon diversity index (p = 0.01) than the mycobiome of ants infected with *Beauveria* (Figure 2B, Supplementary File 2, Table 2). This suggests that the two fungal pathogens investigated, which have different infection strategies, affect the host microbiome and mycobiome differently.

### 3.3 Beta diversity between infection treatments

We also used beta diversity metrics to determine if there were differences in gut micro/mycobiome community compositions between healthy and fungal-infected ants. In doing so, we obtained results that corroborated our alpha diversity results. We again found that bacterial community compositions were not significantly different between infected ants and healthy controls (Figure 2C). However, the fungal community compositions were again significantly different between the *Ophiocordyceps* infected ants and healthy controls (PERMANOVA, p = 0.003) (Figure 2D, Supplementary File 2, Table 3). Moreover, the beta diversity metrics also indicated that both bacterial and fungal community composition differed significantly between ants in the *Ophiocordyceps* and *Beauveria* treatment groups (PERMANOVA, p = 0.033 and p = 0.003, respectively) (Figure 2C and D, Supplementary File 2, Table 3).

In line with findings from previous microbiome work done on *C. floridanus* (25, 68), Enterobacteriaceae (Order Enterobacteriales) constituted the most abundant family of bacteria in the gut microbiome of ants from all treatment groups and were predominately composed of Genus *Blochmannia,* a known endosymbiont of the C*amponotini* tribe. Taxa bar plots for the bacterial microbiome (Figure 2E) show a decrease in the relative abundance of Order Enterobacteriales within the *Beauveria* infection group compared to the *Ophiocordyceps*-infected ants. This might be the underlying cause of the significant difference we observed between the bacterial gut microbiota of *Ophiocordyceps-* and *Beauveria-*infected ants. Taxa bar plots for the gut mycobiome (Figure 2F) showed that *Ophiocordyceps*-infected ants contained a greater percentage of unclassified fungal taxa in their guts compared to the *Beauveria*-infected and healthy control ants, which likely accounts for the found difference in community composition. *Beauveria-*infected and control ants also contained a greater percentage of fungi within Order *Ophiostomatales*, mainly comprised of the species *Sporothrix insectorum,* which is regularly reported to be a fungal pathogen of insects (69, 70).

### 3.4 Differences in alpha diversity in *Ophiocordyceps*-infected ants over time

We compared the alpha diversities of gut microbiota and mycobiota between healthy ants and infected ants at four, progressive time points following *Ophiocordyceps* inoculation. These time points comprised 1) LT25 (lethal time 25%): infected, live ants, before the ultimate manipulated summiting and biting phase; while 25% had succumbed to the infection, 2) LM (live manipulation): live ants that displayed manipulated summiting and biting behavior, 3) DM (dead after manipulation): ants that died a few hours after manipulation took place, and 4) 24PD (24hr post death after manipulation): ants that had been dead for 24 hours after manipulation (Figure 3A and B). Comparisons among these timepoints indicated that observed ASV richness of the bacterial microbiome of LM ants was significantly lower than for healthy controls (p = 0.005), live ants prior to manipulation (LT25, p = 0.022), and those 24 hours after death (24PD, p = 0.03) (Figure 3A) (Supplementary File 2, Table 4). Only the dead manipulated time point (DM), which was taken only a few hours later, was similar to LM. The mycobiome showed the opposite pattern (Figure 3B). Live manipulated (LM) ants had higher fungal ASV richness in their guts than healthy control ants (p = 0.014), infected ants before manipulation (LT25, p = 0.006), and those sampled 24 hours after dying in the manipulated summiting and biting position (24PD, p = 0.048) (Supplementary File 2, Table 4). Alpha diversity differences in the mycobiome among time points were again more apparent than alpha diversity differences in the bacterial microbiome. We observed the same significant differences in Shannon diversity index across timepoints in *Ophiocordyceps-*infected ants (Figure 3B); Live manipulation samples had a higher Shannon diversity index compared to healthy control samples (p < 0.0001), LT25 (p = 0.012), and 24PD samples (p = 0.034) (Supplementary File 2, Table 4). Taken together, these results suggest that the gut micro- and mycobiome of ants that show *Ophiocordyceps*-adaptive summiting and biting behaviors are significantly different compared to healthy ants, and even those in other stages of the infection.

### 3.5 Beta diversity between *Ophiocordyceps*-infected ants over time

To investigate potential differences in gut microbiome and mycobiome composition between behaviorally manipulated ants (LM) and other stages of infection, we used a weighted beta diversity metric (Bray-Curtis) and relative abundances of community constituents. For the bacterial microbiome of *Ophiocordyceps-*infected ants (Figure 3C), we only found a significant difference in community composition between the guts of live manipulated ants (LM) and those that were sampled 24 hours after manipulation (24PD) and death (DM) (PERMANOVA, p = 0.04) (Supplementary File 2, Table 5). In contrast, the mycobiome of *Ophiocordyceps-*infected ants (Figure 3D) showed significant differences in community composition for several timepoint comparisons. The gut mycobiome of ants that were sampled during manipulated summiting behavior (LM) had significantly different fungal community composition than healthy control ants (p = 0.01), infected ants prior to manipulation (LT25, p = 0.02) and ants that were sampled 24 hours after they succumbed to the infection (24PD, p = 0.01) (Supplementary File 2, Table 5). These results support the evidence from the alpha diversity analyses that the fungal mycobiome is significantly different in *Ophiocordyceps*-infected ants during manipulated summiting and biting behavior as compared to in healthy ants and at other infection time points. Taxa bar plots (Figure 3F) indicate a sudden increase in the relative abundance of unclassified taxa at the behavioral manipulation (LM) timepoint and a large decrease in the relative abundance of Order *Ophiostomatales,* the most prevalent fungal order observed in our samples. In the samples from the two time points taken after behavioral manipulation (DM and 24PD), there is again an increase in the relative abundance of *Ophiostomatales* and a decrease in unclassified taxa, indicating a gradual shift in community composition.

### 3.6 Alpha and beta diversity in *Beauveria*-infected ants over time

We also looked into the gut micro- and mycobiota of ants infected with *Beauveria* at infection time points that we consider to parallel those investigated for *Ophiocordyceps* infection. As such, we sampled *Beauveria*-infected ants at LT25, when showing non-manipulative “sick behavior” (LS), having died after showing sick behavior (DS) and 24 hours post death (24PD). While the bacterial and fungal gut alpha diversities of *Ophiocordyceps*-infected ants that showed the manipulation phenotype (LM) were different from healthy and other infection stages, the gut composition of *Beauveria*-infected ants that showed the paralleling “sick” phenotype (LS) was not. Comparing bacterial and fungal alpha diversities across infection time points, we only detected a significant difference in the observed richness of the fungal mycobiome between healthy control ants and *Beauveria*-infected ants that had not yet shown the sick phenotype (LT25, p = 0.034) (Supplementary File 2, Table 6). However, the results for analyses of Shannon diversity did not corroborate this finding (Figure 4B). Additionally, our beta diversity analyses gave another, singular and distinct result. Using the Bray-Curtis metric, we found that the 24-hour postmortem time point (24PD) was the only time point at which the bacterial microbiome composition was significantly different between *Beauveria*-infected and healthy control ants (PERMANOVA, p=0.01). The bacterial microbiome composition also differed significantly between the early *Beauveria*-infection time point in which a sick phenotype was not detected yet (LT25) and the 24PD time point (PERMANOVA, p = 0.04) (Figure 4C, Supplementary File 2, Table 7). These results suggest that *Beauveria* infection does not affect the host bacterial microbiome or fungal mycobiome consistently enough to detect significant differences between infection stages. Moreover, the taxa bar plots from different *Beauveria* infection time points indicated that the mycobiome of *Beauveria-*infected ants (Figure 4F) was dominated by *Ophiostomatales* while the relative abundance of unidentified fungal taxa was a lot smaller. This is in stark contrast with the relative abundances of fungal gut taxa that we observed in the guts of *Ophiocordyceps*-infected ants. There, we found and relative increase of unidentified fungal gut taxa, while the abundance of *Ophiostomatales* decreased. The increased relative abundance of these unidentified fungal gut taxa, especially during behavioral manipulation, might be a specific hallmark of the infection strategies of *Ophiocordyceps*.

## 4. Discussion

Fungal pathogens can have vastly different infection strategies, ranging from necrotrophic to biotrophic. In addition to affecting the host and its tissues differently, these pathogens may impact the host’s microbiota in different ways as well. To test this hypothesis for insect-infecting fungi specifically, we investigated the effects of infection with *O. camponoti-floridani* and *B. bassiana* on the gut microbiota and mycobiota of the carpenter ant (*C. floridanus*). *Ophiocordyceps camponoti-floridani* is a specialist, behavior-manipulating, hemi-biotroph-like pathogen of *C. floridanus*, and *B. bassiana* is a generalist, non-manipulating, necrotroph-like pathogen. When comparing the bacterial microbiome (16S) of healthy ant guts to their *Ophiocordyceps*- and *Beauveria*-infected counterparts we did not find any significant differences. The bacterial community composition of infected ants remained relatively similar to that of healthy controls. This would indicate that fungal infection is not impacting the ant gut microbiome as hypothesized. However, we found that the fungal mycobiome (ITS) of *Ophiocordyceps-*infected ants had a significantly higher ASV diversity (Shannon index) than the mycobiome of healthy control ants with corroborating differences in community composition. Thus, based on the results we observed for gut mycobiota, we conclude that behavior-manipulating *Ophiocordyceps* infection does, in fact, affect the host gut. During *Ophiocordyceps* infection, the fungal mycobiome appears to be more dynamic in its composition as compared to the bacterial microbiome, with relative abundances of fungal gut taxa changing upon infection. *Blochmannia* was by far the most dominant bacterial genus within the carpenter ant microbiome, both in this and other studies (25, 68), which may have obscured changes other than those in the dominating species. Instead, there may be higher evenness between fungal taxa than between bacterial taxa, making subtle differences in the mycobiome easier to detect. Studies in other insect species also suggest that the fungal mycobiome may be more variable than the bacterial microbiome (36, 71). Moreover, we detected a dominant presence of *S. insectorum*, which is commonly known as an entomopathogen (69, 70), but not necessarily as an insect gut symbiont. In line with this, other known entomopathogens, of the genus *Ophiocordyceps,* have been detected in the mycobiomes of plants and beetles (72, 73). As such, exploring the mycobiome can also reveal alternative lifestyles of presumed primary pathogens. Moreover, studies in humans and other animals (including disease vectors such as mosquitoes (74)) clearly indicate that the fungal mycobiome influences disease susceptibility and progression (reviewed in (75, 76)). Taken together, this all suggests that it is worthwhile to include the mycobiome in insect microbiota studies, which are currently still largely overlooked.

We also found that both bacterial and fungal diversity and composition of the gut community were significantly different between ants infected with *Ophiocordyceps* and *Beauveria*. The bacterial microbiome of *Ophiocordyceps*-infected ants had a significantly lower ASV diversity than the microbiome of the *Beauveria*-infected ants. In contrast, the mycobiome of *Ophiocordyceps*-infected ants had a significantly higher ASV diversity than in *Beauveria*-infected ants. These results indicate that these entomopathogenic fungi, with contrasting infection strategies, impact the overall gut microbiota differently. Additionally, the specialist, behavior-manipulating fungus *Ophiocordyceps* caused a significant dysbiosis while the generalist fungus *Beauveria* did not seem to affect the gut composition much.

Indeed, when we compared healthy control ants to ants at progressive infection time points, we did not detect consistent, significant changes in the gut communities of *Beauveria*-infected ants when analyzing alpha and beta diversity. This suggests that *Beauveria* infection leaves the gut composition relatively unchanged, with perhaps some gradual changes leading up to the fungus’ saprophytic growth after death as the significant differences in beta diversity between the 24 hours post death (24PD) time point and the healthy and early infection time point (LT25) suggest.

In contrast, in *Ophiocordyceps*-infected ants, the micro- and mycobiome both displayed a notable shift in ASV richness at live manipulation (LM) relative to both earlier (i.e., healthy control and LT25) and later (i.e., 24PD) infection time points. During this time, the ant host shows a characteristic summiting behavior adaptive to the development and spread of the fungal pathogen, *Ophiocordyceps* (9, 10). Alpha diversity for both micro- and mycobiomes differed significantly between ants at the LM time point and ants from the healthy control group, and the LT25 and 24PD timepoints, which may be an indication of a gradual change with infection progression. However, alpha diversity of the micro- and mycobiome did not differ between the other time points, suggesting that something uniquely different might be happening with gut microorganisms during manipulated summiting behavior. At the LM timepoint, the microbiome again showed a sudden drop in bacterial ASV richness while there was a significant increase in fungal ASV richness seen for the mycobiome. The only time point during which the gut micro- and mycobiota was not significantly different compared to the LM timepoint was death after manipulation (DM). For *Ophiocordyceps*-infected ants, death occurs within a mere few hours of live manipulation. In contrast, the other time points were separated from the LM time point by 24+ hours (24PD) to several days (LT25). As such, the relatively few hours difference between LM and DM time points may not have been enough time for the micro- and mycobiota to change significantly. However, the micro- and mycobiota of LM ants was significantly different from that of 24PD ants, while we did not observe significant differentiation between the DM and 24PD timepoints. This suggests that there is already a gradual change in ant gut micro- and mycobiota in the short amount of time between being live manipulated and eventually succumbing to the infection. While the beta diversity analysis corroborated these findings for the mycobiome, they did not do so entirely for the bacterial microbiome. This suggests again that the fungal composition of the gut appears to be more dynamic than the bacterial composition and that *Ophiocordyceps* infection is somehow significantly affecting that composition, especially during the manipulated summiting stage.

These different disease outcomes for the host gut micro- and mycobiota could be due to a direct effect of the pathogen on the host and its tissues. Alternatively, it could be a result of changed behavior upon infection. Many social insects, including *Camponotus* ants, engage in trophallaxis – the social exchange of fluids between individuals. Such social fluids contain partially digested and regurgitated food materials as well as that individual’s endocrine molecules that play a role in maintaining the social network and decision-making within an ant colony. Trophallaxis is also hypothesized to play a role in inoculating and maintaining gut microbiota among colony members (77). Thus, a disconnect from the social network of the colony could have potential impacts on an individual’s microbiome. Indeed, *Ophiocordyceps*-infected ants display diminished communication with nestmates (13), which consequently might reduce trophallaxis. As such, they might be disconnected from their colony mates with the consequence that their regular gut microbiota is not maintained. While the period between infection, manipulation and eventual death spans several weeks in *Ophiocordyceps, Beauveria* infection runs a more rapid course of 5 days to a week. Diminished communication was not readily observed in *C. floridanus* infected with *Beauveria* (13). However, even if reduced communication were to occur, the disruption might be too brief to have a measurable influence on the micro- and mycobiota. Therefore, a prolonged detachment from the social-fluid exchange due to asocial behavior might result in the microbiome dysbiosis seen in *Ophiocordyceps*-infected ants, which could explain the differences between the microbiomes of *Ophiocordyceps*-versus *Beauveria*-infected ants. However, if this were the case, one might expect a more gradual change over time with significance compared to healthy controls being found for all later stages of infection (LM, DM and 24PD) and not just the single live manipulation time point.

Of course, we cannot completely exclude that our non-significant results are caused by the nature of our samples and sample size. The survival curve for the *Ophiocordyceps* infections (Figure 1A) followed expected trends with most infected ants dying once manipulations began (21 DPI). The survival curve for *Beauveria* infections (Figure 1B) also matched expected trends with a decline in survival starting between 3-5 DPI. However, the survival rate levelled out after 6 DPI with around 50% of the population still alive. This indicates that not all ants progressed through the infection the same way and that some were able to clear it. The sick (LS), dead (DS), and 24 post death (24PD) *Beauveria* timepoint samples are unaffected by this, as they were sampled based on a distinguishable sick behavior phenotype. However, the LT25 timepoint samples may be a mixture of infected and uninfected individuals as there was no clear way to distinguish infection status for those ants. Moreover, sample size was limited for each timepoint and might not have been robust enough to reveal the full effects of fungal infection over time. Yet, the presence of dysbiosis in the guts of *Ophiocordyceps*-infected ants that displayed summiting behavior was clearly significantly different compared to most other infection time points, including healthy ants. In fact, looking at the NMDS plots (Figure 3C and D), the LM samples cluster together while the other biological groups seem to show a more dispersed level of variation. As such, we consider the finding that *Ophiocordyceps* infection affects the gut micro- and mycobiome of ants during manipulated summiting to be evident.

The biological relevance of a distinctly altered gut microbiome composition during manipulated summiting behavior remains a topic of speculation. One possibility could be that the ant gut microbiome affects host behavior and may be involved, to some degree, in the expression of stereotypically altered behaviors during *Ophiocordyceps* infection. There is mounting evidence from human research that gut dysbiosis is associated with an increased risk of Alzheimer’s, depression, and other behavioral pathologies (78). Some studies suggest that this may occur via microbial metabolites secreted in the gut interacting with neurological receptors and G protein coupled receptors (GPCRs) (79, 80). Transcriptomic analyses of *Ophiocordyceps*-infected ants that displayed manipulated summiting behavior showed enrichment for GPCRs among the genes that were downregulated during this infection stage as compared to healthy individuals (8). These GPCRs were annotated to have functions involved in the perception of light and smell, and the binding of biogenic amines such as dopamine. Most of the pathogen-related behavioral changes modulated by these GPCRs are likely due to the predicted binding of *Ophiocordyceps* secreted proteins that are upregulated during manipulated summiting (20). However, other GPCRs might be interacting with (altered levels of) microbial peptides secreted by the microbiome. Additionally, there is increasing evidence that gut microbiomes aid in insect host immune defense and microbiome homeostasis is thought to be an important factor in mediating the immune system (26, 81). For instance, recent research into honeybee core microbiomes showed that dysbiosis was linked to higher susceptibility to pathogen invasion and consequently higher mortality rates (81). Considering that a hemibiotroph-like pathogen such as *Ophiocordyceps* needs to ensure prolonged colonization and successful manipulation of the ant without causing significant physical damage, this fungus could perhaps be affecting the microbiome such that the host immune response is effectively lowered. This hypothesis would be in line with metabolomics data that suggest a lowered immune response in the ant paired with the presence of neuroprotectants (19, 82). However, one would expect this strategy to be needed during the earlier time points in infection as well, where we did not detect any significant micro-nor mycobiome changes. Taken together, the potential pathogen-adaptive function of gut dysbiosis during *Ophiocordyceps* infection, if any, remains to be discovered. Nevertheless, both the hypotheses we propose here could provide an explanation as to why we do detect this dysbiosis of the host gut in infections with the hemibiotroph-like, behavior-manipulating specialist *Ophiocordyceps* and not in those with the necrotroph-like, non-manipulating generalist *Beauveria*.

## 5. Conclusions

To our knowledge, all previous research done on the gut microbiota of ants has focused on their bacterial microbiome, making our study the first to investigate the fungal taxa present within the ant gut, the mycobiome. Our data show that, while the bacterial microbiome is often dominated by one species, the fungal mycobiome is highly diverse with more evenness between taxa. Moreover, our data suggest that the fungal taxonomic composition of the ant gut is more dynamic, as community changes were more prominently detected in the fungal components of the gut. These results support the notion that the fungal portion of the microbiome likely plays a more prominent role in insect functioning than previously credited and should be studied more intensely. Moreover, we provide the first evidence that the gut microbiome of *C. floridanus*, and with that perhaps also other insects, appears to be impacted differently by infecting entomopathogens with different lifestyles. More specifically, our data suggest that behavior-manipulating specialists like *O. camponoti-floridani* might affect the host microbiome in a timed manner that aligns with pathogen-adaptive host behavioral phenotypes. As such, considering the role that microbiota might play in host behavior, part of the behavioral manipulation might be the result of dysbiosis of the gut microbiota. Research in which specific modulation of gut taxa is linked to host behavior and infection status will be needed to begin to unravel these potential intersections. This will improve our understanding of the gut-brain axis and provide knowledge that could improve the effective use of fungal pathogens in the biocontrol of insect pests.

## CRediT authorship contribution statement

**Sophia Vermeulen:** Data curation, Formal Analysis, Investigation, Methodology, Writing – original draft. **Anna Forsman:** Formal Analysis, Methodology, Supervision, Writing – review and editing. **Charissa de Bekker:** Conceptualization, Formal Analysis, Funding acquisition, Methodology, Supervision, Writing – review and editing.

## Acknowledgements

We would like to thank Ian Will, Biplabendu Das, and Erandeni Ponce de Leon for their assistance with ant collection, infection, and monitoring. We would also like to thank the Fall 2022 UCF Genomics Course undergraduate students for being involved with the DNA extractions. Funding to conduct this study was obtained from the National Science Foundation, CAREER IOS-1941546, awarded to Charissa de Bekker.

## Appendix. Supplementary materials

**Supplementary File 1:** This file presents figures and tables that support our technical data and sample validation analyses.

**Supplementary File 2:** This file contains tables that present the output of the alpha and beta diversity statistical analyses performed in this study.

## Data availability

All sequence data generated in this study have been deposited in the NCBI Sequence Read Archive under BioProject ID PRJNA1129269.

## Supplementary

**Supplementary Figure 1:**
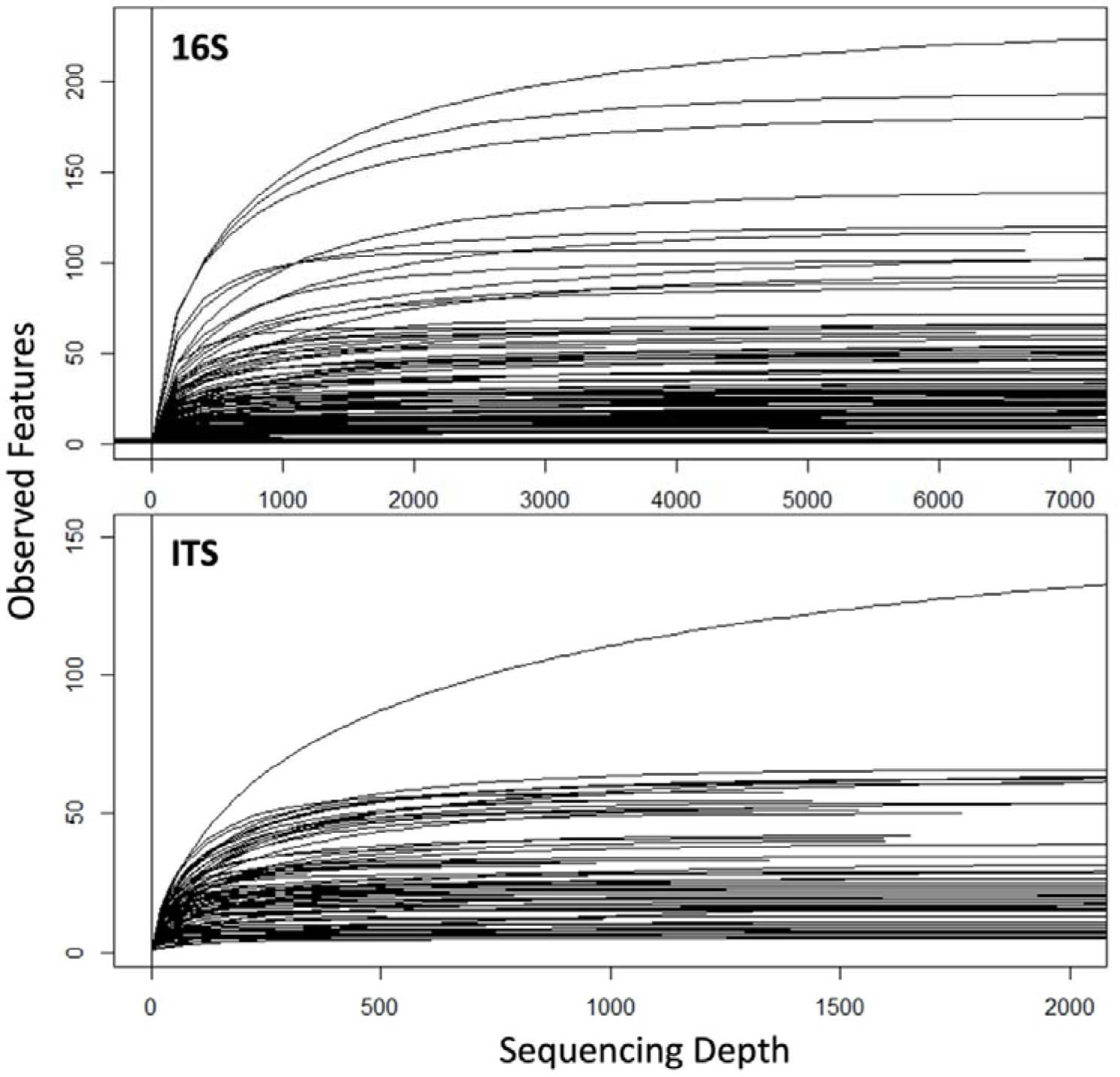
Rarefaction Curves for 16S (top) and ITS reads (bottom). Rarefaction curves indicate sufficient species coverage at a sequence sampling depth of 4000 for 16S libraries, and 500 for ITS libraries. Observed features represent the number of different ASV’s identified.

**Supplementary Table 1:** Samples remaining after read filtering and rarefaction. Ten samples were collected at each timepoint, except for Ophiocordyceps ‘live manipulated’ where 13 samples were collected and sequenced. Control samples were pooled for both experiments, from which we randomly selected 24 samples for processing.

|  | 16S libraries |  |  | ITS libraries |  |  |
| --- | --- | --- | --- | --- | --- | --- |
|  | *Ophiocordyceps* | *Beauveria* | Control | *Ophiocordyceps* | *Beauveria* | Control |
| **Total** | **33** | **37** | **24** | **38** | **23** | **20** |
| LT25 | 8 | 10 |  | 8 | 7 |  |
| LM/LS | 11 | 10 |  | 13 | 5 |  |
| DM/DS | 9 | 9 |  | 8 | 6 |  |
| 24PD | 5 | 8 |  | 9 | 5 |  |

**Supplementary Figure 2:**
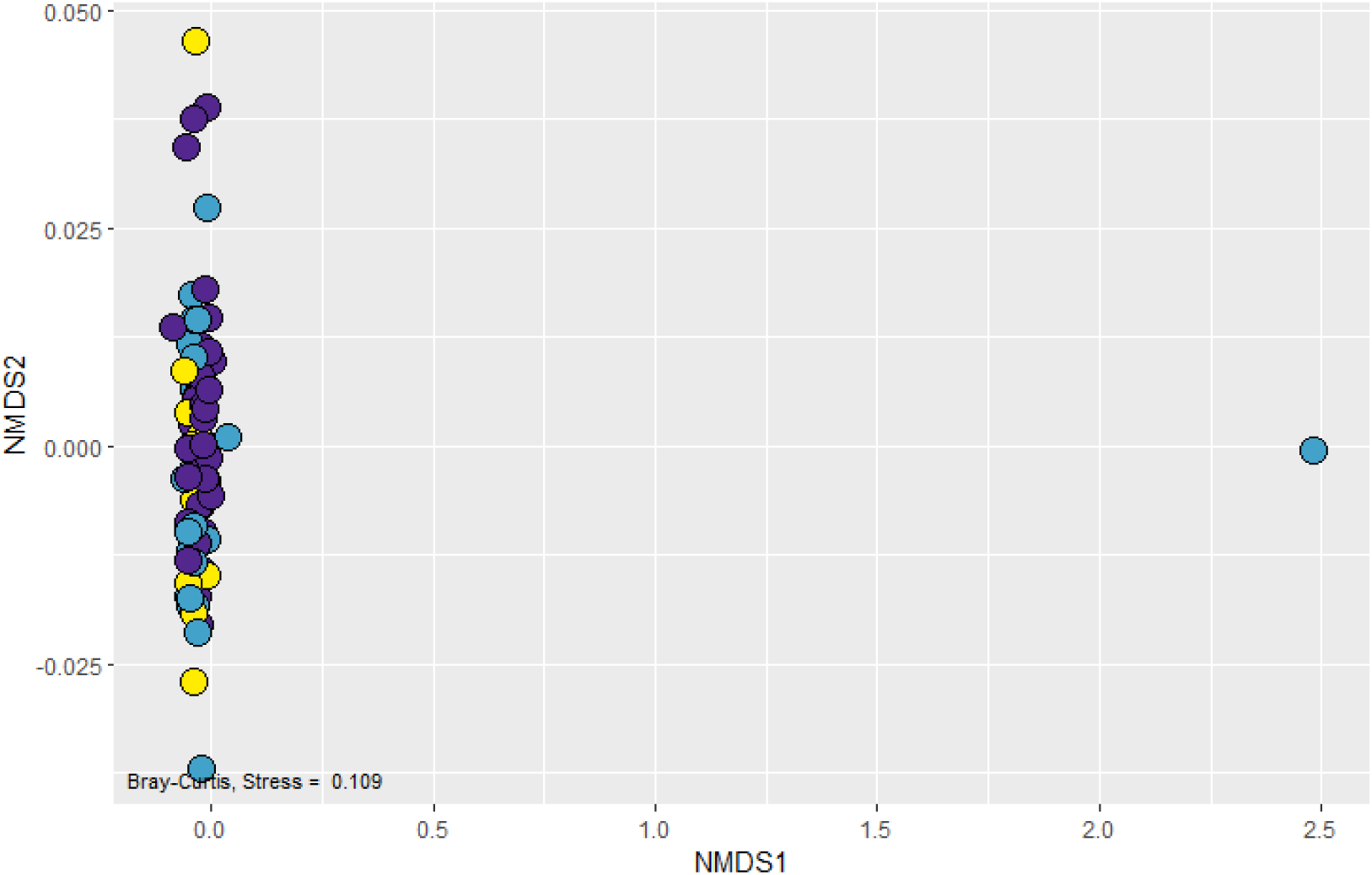
ITS NMDS plot with outlier Beauveria sample included. Non-metric multidimensional scaling (NMDS) plot using a Bray Curtis distance matrix (stress = 0.109) of the gut mycobiome (ITS) for different treatment groups. The points represent microbiomes of individual samples, and the colors indicate which treatment groups the samples belong to (yellow = control, purple = Ophiocordyceps, blue = Beauveria). The blue dot at the far right of the graph represents the outlier from the Beauveria treatment group that was removed for analysis. After applying read filtering, and removal of Ophiocordyceps and Beauveria reads, this sample was only left with unidentified reads and reads identified as Malassezia, a likely contaminant.

**Supplementary Figure 3:**
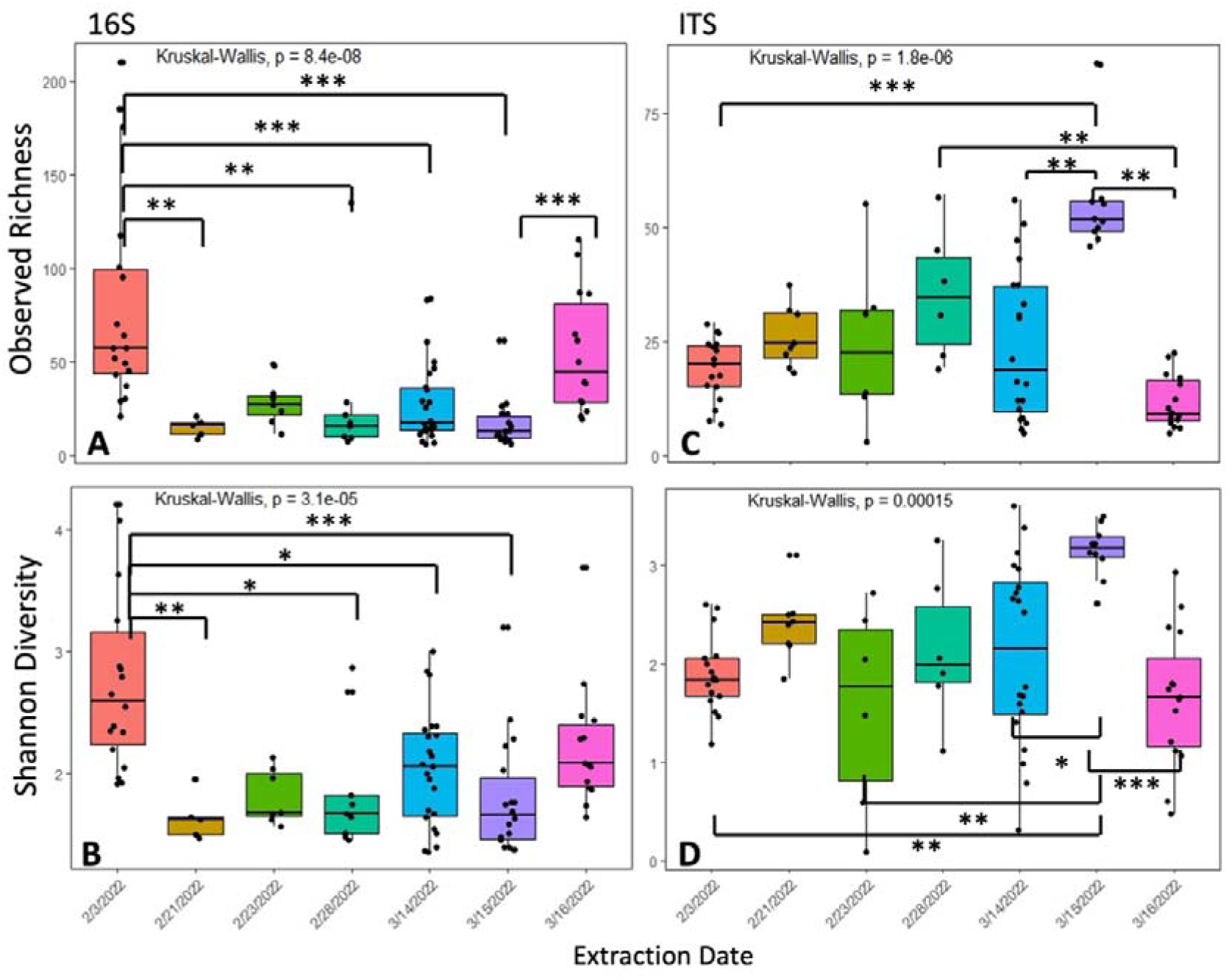
Alpha diversity plots of extraction batches for 16S and ITS libraries. A) Observed richness in the bacterial microbiome (16S) of extraction batches containing random samples. B) Shannon diversity index of the bacterial microbiome of extraction batches. C) Observed richness for the mycobiome (ITS) of extraction batches. D) Shannon diversity index for the mycobiome of extraction batches. P-value significance between extraction batches containing random samples from this study: * <0.05, ** < 0.01, *** <0.001.

**Supplementary Figure 4:**
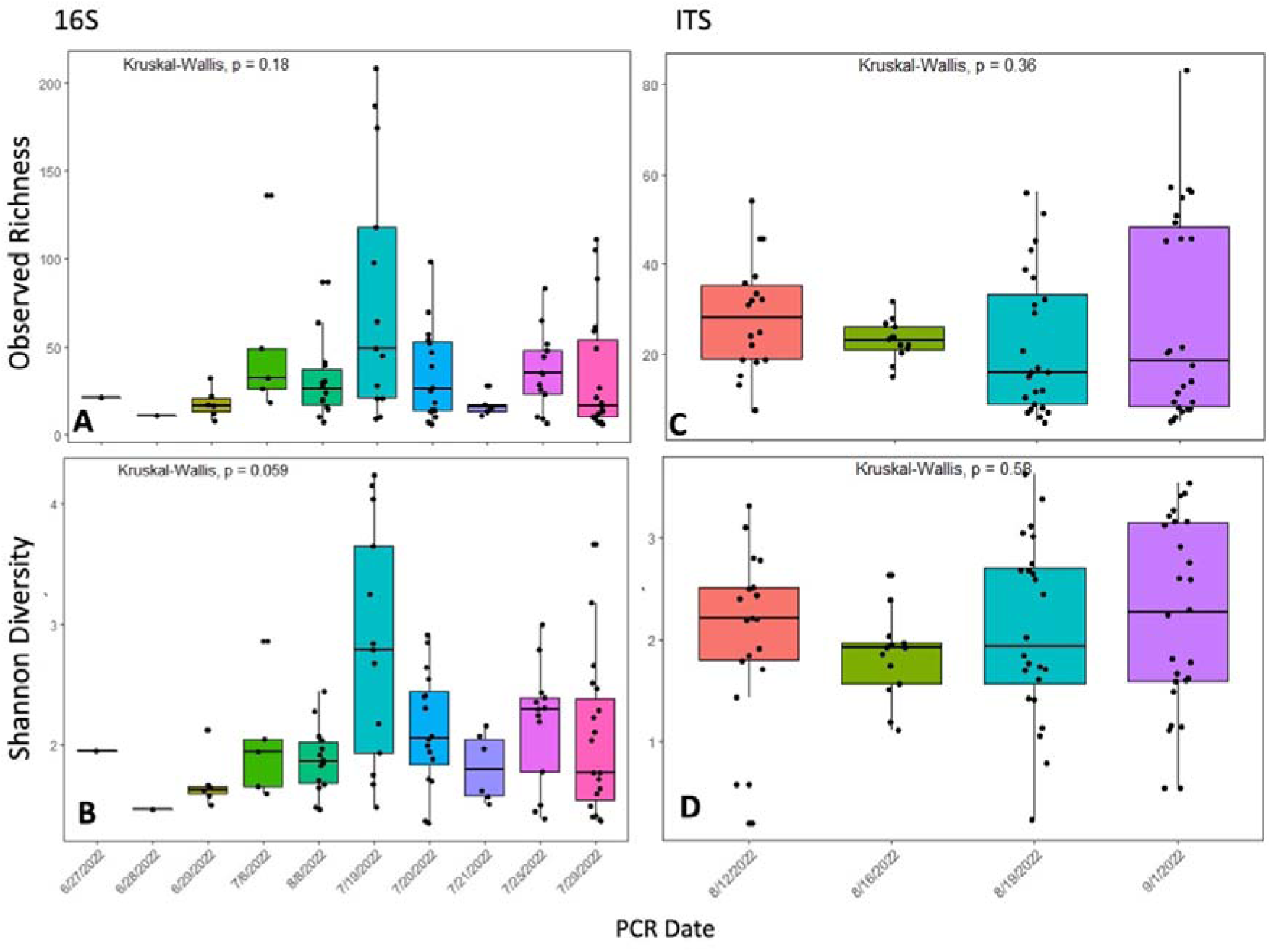
Alpha diversity plots of PCR batches for 16S and ITS libraries. A) Observed richness in the bacterial microbiome (16S) of PCR batches containing random samples from this study. B) Shannon diversity index of the bacterial microbiome of PCR batches. C) Observed richness for the mycobiome (ITS) of PCR batches. D) Shannon diversity index for the mycobiome of PCR batches. No significant differences were seen between PCR batches containing random samples from this study.

**Supplementary Table 2:** Dunn test results for alpha diversity comparisons between treatments. P-values have been adjusted with the Bonferroni correction for multiple comparisons. P-values in bold font indicate significance (p < 0.05).

|  | Observed Richness |  | Shannon Diversity |  |
| --- | --- | --- | --- | --- |
| Pairwise comparison | 16S | ITS | 16S | ITS |
| *Ophiocordyceps* - *Beauveria* | **0.003** | **0.006** | **0.0003** | **0.01** |
| *Ophiocordyceps* - Control | 0.123 | 0.69 | 0.63 | **0.01** |
| *Beauveria* - Control | 1 | 0.34 | 0.073 | 1 |

**Supplementary Table 3:** Pairwise PERMANOVA results for the beta diversity comparisons between treatments. P-values have been adjusted with the Bonferroni correction for multiple comparisons. P-values in bold indicate significance (p<0.05).

| Pairwise comparison | Df | SumOfSqs | F.Model | R2 | P-value |
| --- | --- | --- | --- | --- | --- |
| *Ophiocordyceps* - *Beauveria* | 1 | 0.368 | 3.04 | 0.043 | **0.033** |
| *Ophiocordyceps* - Control | 1 | 0.098 | 1.18 | 0.021 | 0.882 |
| *Beauveria* – Control | 1 | 0.320 | 2.47 | 0.040 | 0.069 |

| Pairwise comparison | Df | SumOfSqs | F.Model | R2 | P-value |
| --- | --- | --- | --- | --- | --- |
| *Ophiocordyceps* – *Beauveria* | 1 | 1.6 | 4.61 | 0.0725 | **0.003** |
| *Ophiocordyceps* - Control | 1 | 1.58 | 4.84 | 0.0795 | **0.003** |
| *Beauveria* - Control | 1 | 0.34 | 1.03 | 0.0245 | 1 |

**Supplementary Table 4:**
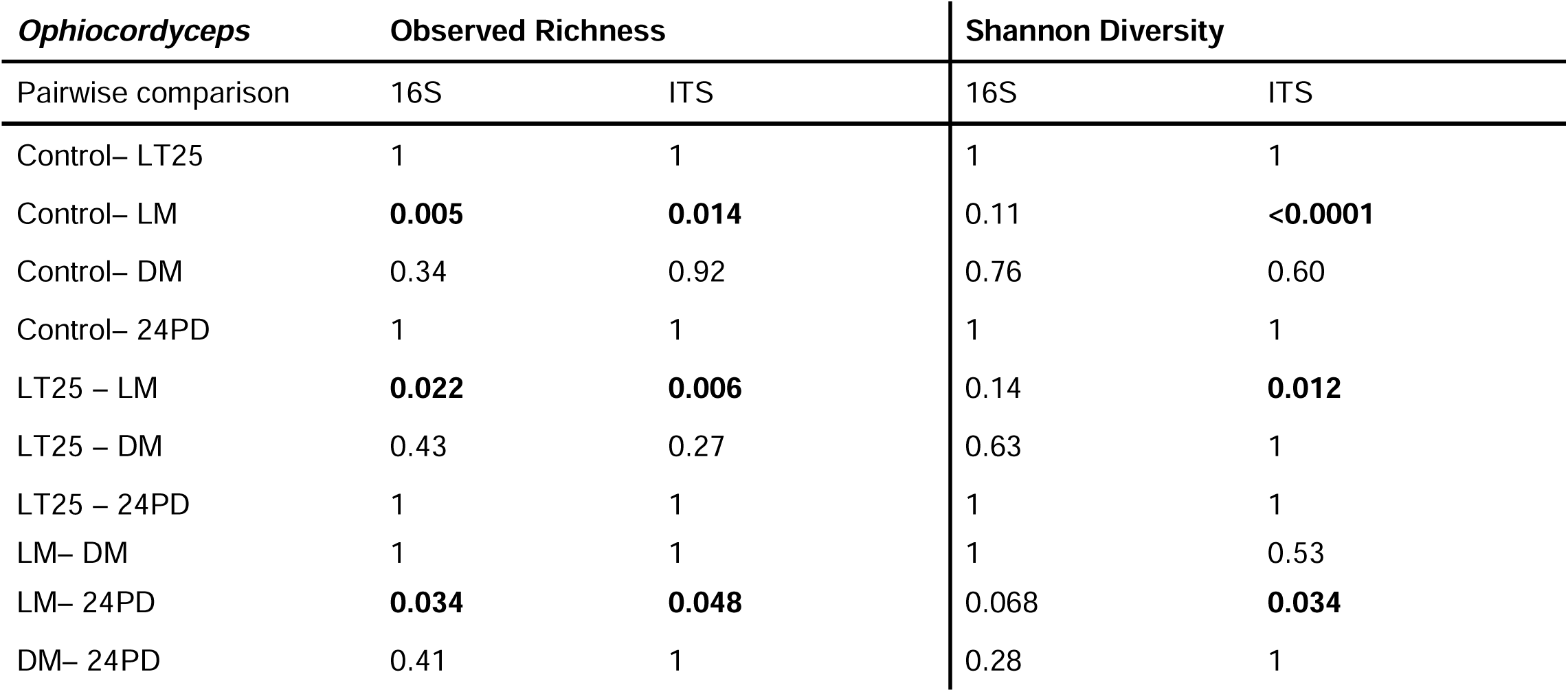
Dunn test results for alpha diversity comparisons between Ophiocordyceps infection timepoints. P-values have been adjusted with the Bonferroni correction for multiple comparisons. P-values in bold font indicate significance (p < 0.05).

**Supplementary Table 5:**
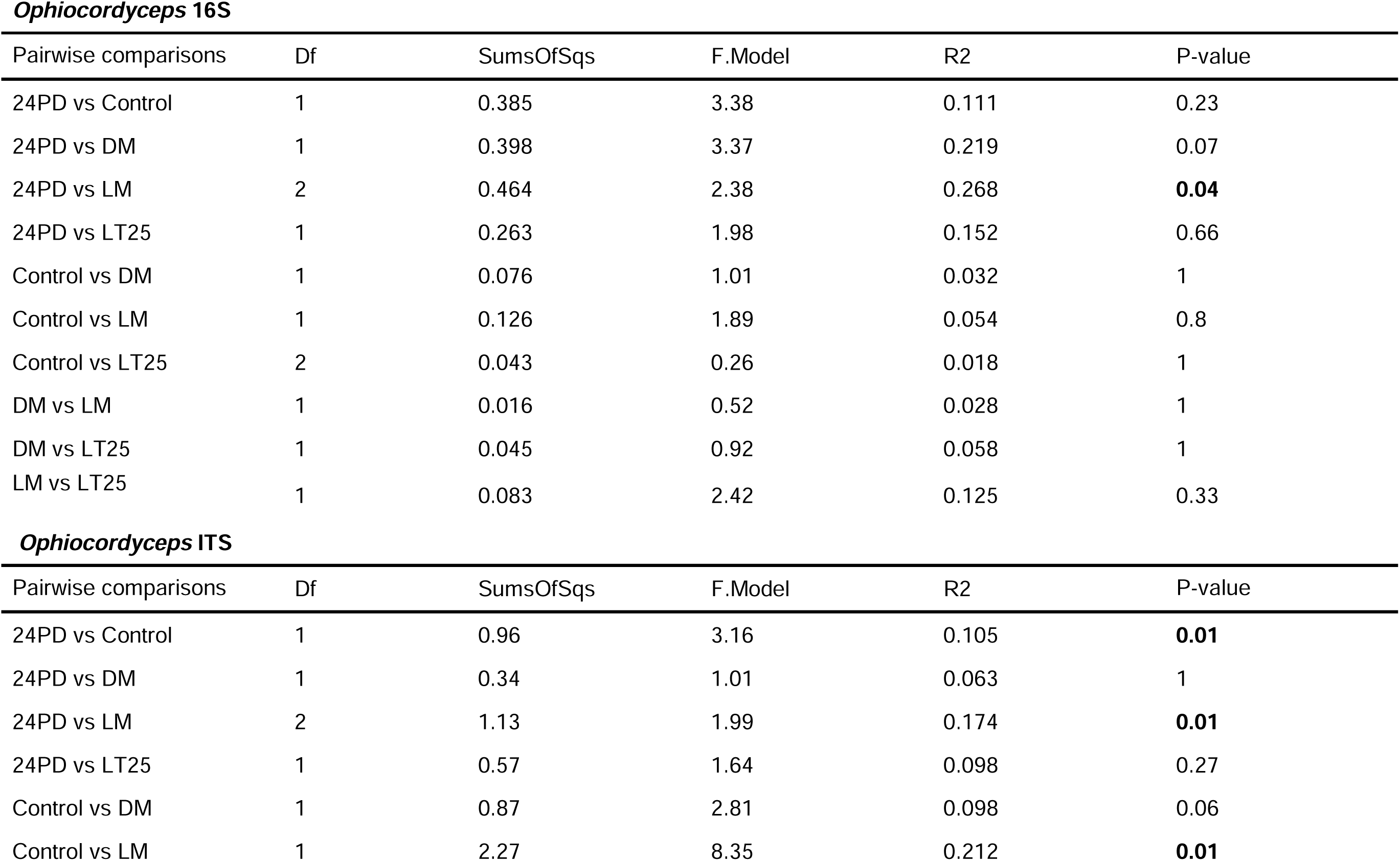
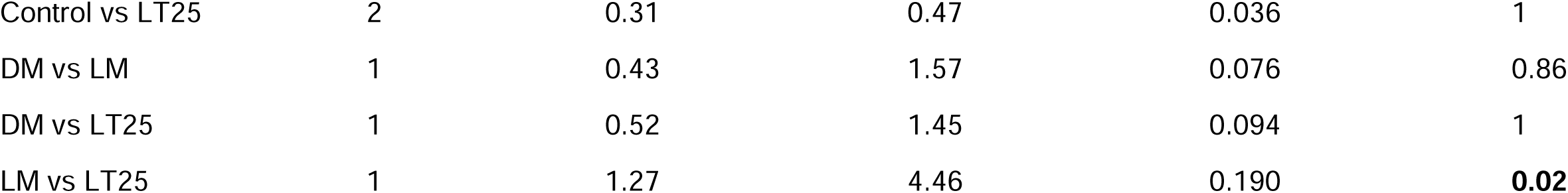
Pairwise PERMANOVA results for the beta diversity comparisons between Ophiocordyceps infection timepoints. . P-values have been adjusted with the Bonferroni correction for multiple comparisons. P-values in bold indicate significance (p<0.05).

| Pairwise comparisons | Df | SumsOfSqs | F.Model | R2 | P-value |
| --- | --- | --- | --- | --- | --- |
| 24PD vs Control | 1 | 0.385 | 3.38 | 0.111 | 0.23 |
| 24PD vs DM | 1 | 0.398 | 3.37 | 0.219 | 0.07 |
| 24PD vs LM | 2 | 0.464 | 2.38 | 0.268 | **0.04** |
| 24PD vs LT25 | 1 | 0.263 | 1.98 | 0.152 | 0.66 |
| Control vs DM | 1 | 0.076 | 1.01 | 0.032 | 1 |
| Control vs LM | 1 | 0.126 | 1.89 | 0.054 | 0.8 |
| Control vs LT25 | 2 | 0.043 | 0.26 | 0.018 | 1 |
| DM vs LM | 1 | 0.016 | 0.52 | 0.028 | 1 |
| DM vs LT25 | 1 | 0.045 | 0.92 | 0.058 | 1 |
| LM vs LT25 | 1 | 0.083 | 2.42 | 0.125 | 0.33 |

***Ophiocordyceps* ITS**
| Pairwise comparisons | Df | SumsOfSqs | F.Model | R2 | P-value |
| --- | --- | --- | --- | --- | --- |
| 24PD vs Control | 1 | 0.96 | 3.16 | 0.105 | **0.01** |
| 24PD vs DM | 1 | 0.34 | 1.01 | 0.063 | 1 |
| 24PD vs LM | 2 | 1.13 | 1.99 | 0.174 | **0.01** |
| 24PD vs LT25 | 1 | 0.57 | 1.64 | 0.098 | 0.27 |
| Control vs DM | 1 | 0.87 | 2.81 | 0.098 | 0.06 |
| Control vs LM | 1 | 2.27 | 8.35 | 0.212 | **0.01** |

|  |  |  |  |  |  |
| --- | --- | --- | --- | --- | --- |
| Control vs LT25 | 2 | 0.31 | 0.47 | 0.036 | 1 |
| DM vs LM | 1 | 0.43 | 1.57 | 0.076 | 0.86 |
| DM vs LT25 | 1 | 0.52 | 1.45 | 0.094 | 1 |
| LM vs LT25 | 1 | 1.27 | 4.46 | 0.190 | **0.02** |

**Supplementary Table 6:**
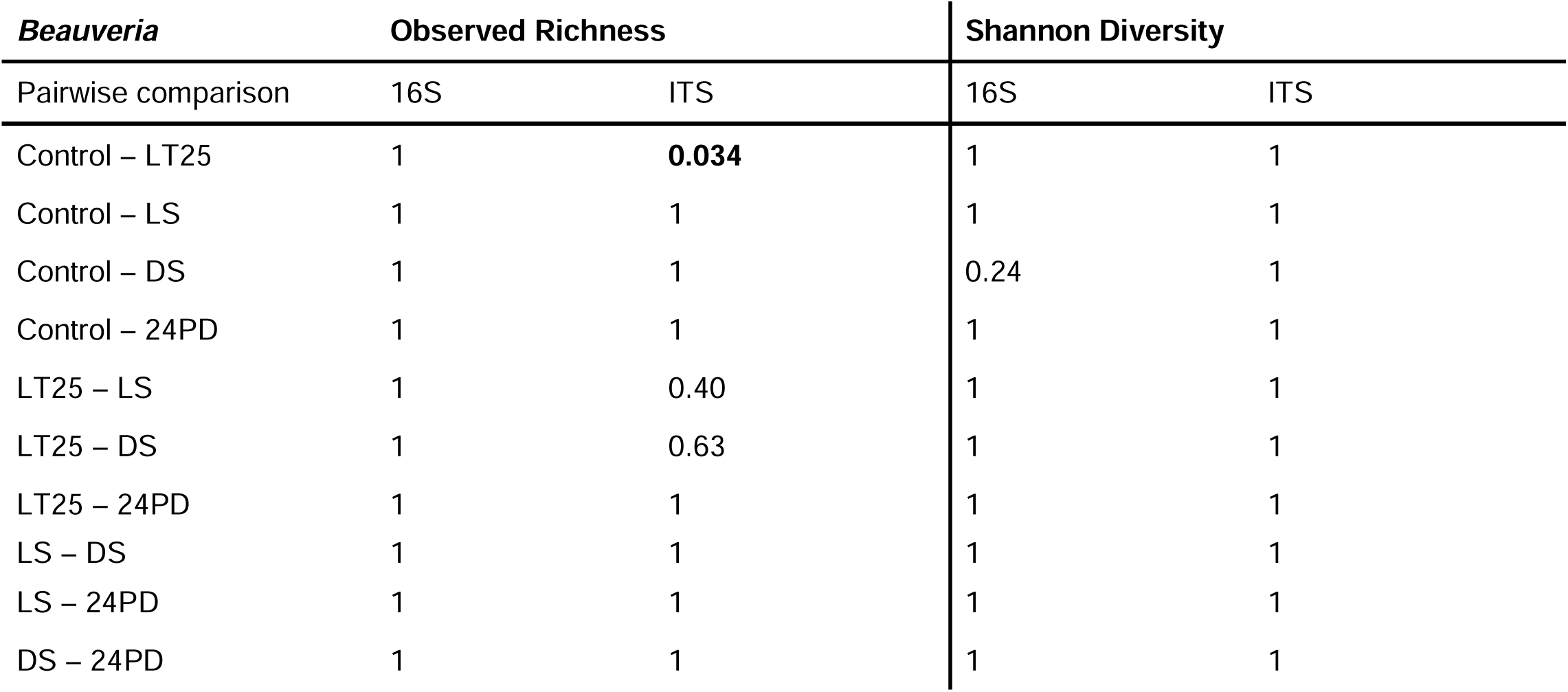
Dunn test results for alpha diversity comparisons between Beauveria infection timepoints. P-values have been adjusted with the Bonferroni correction for multiple comparisons. P-values in bold font indicate significance (p < 0.05).

**Supplementary Table 7:**
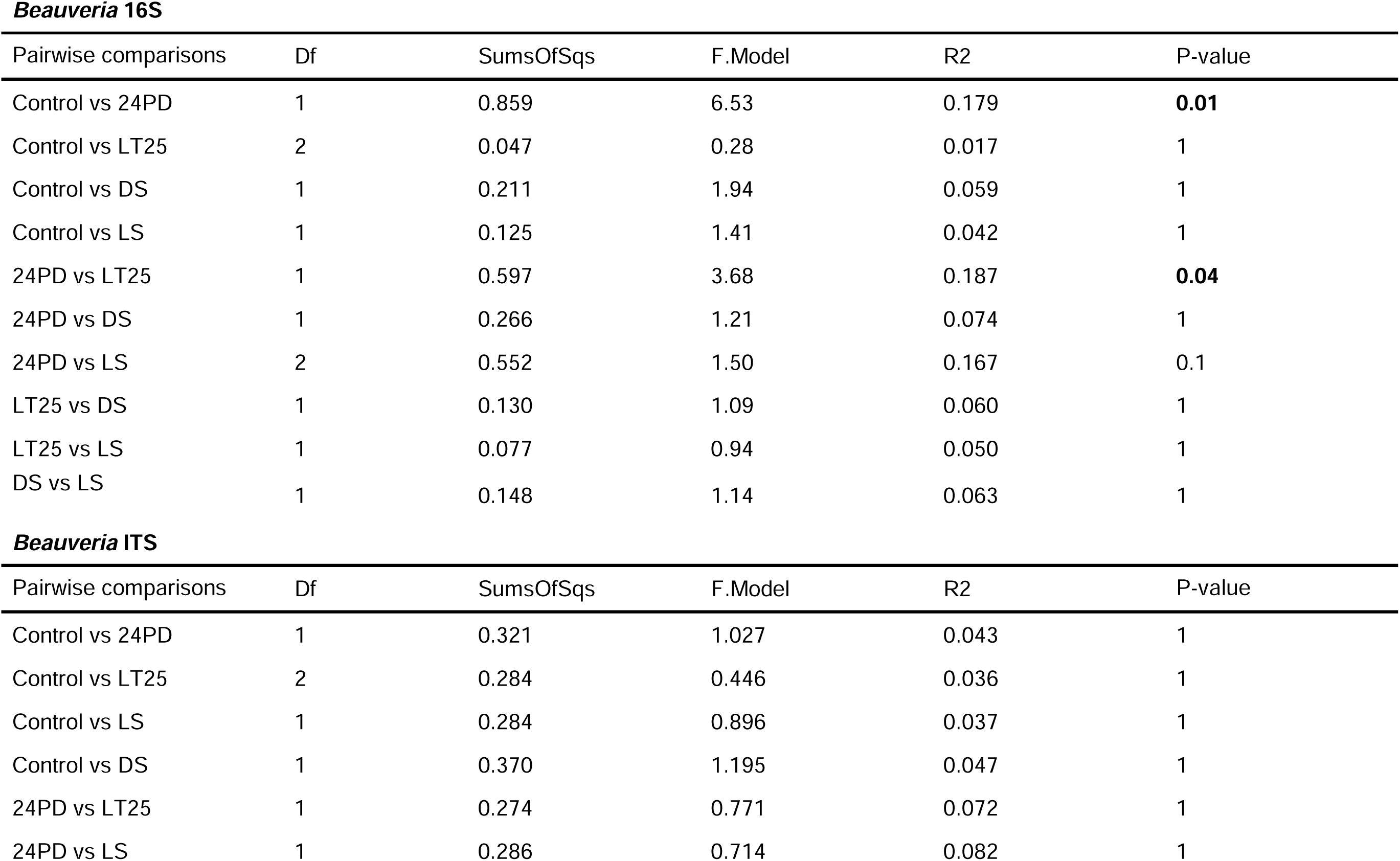
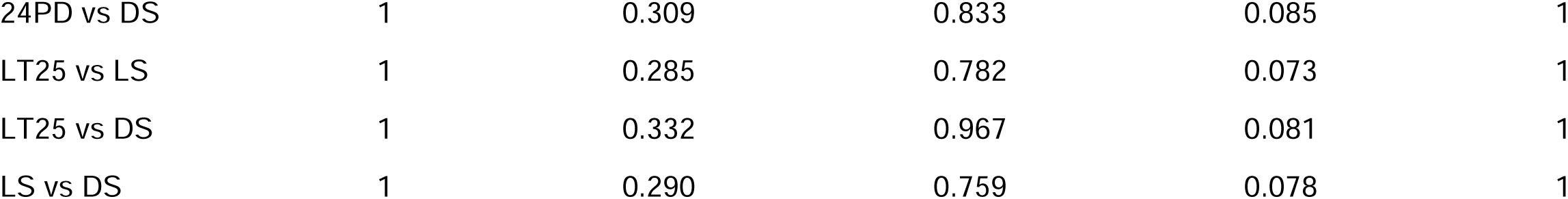
Pairwise PERMANOVA results for the beta diversity comparisons between Beauveria infection timepoints. P-values have been adjusted with the Bonferroni correction for multiple comparisons. P-values in bold indicate significance (p<0.05).

| Pairwise comparisons | Df | SumsOfSqs | F.Model | R2 | P-value |
| --- | --- | --- | --- | --- | --- |
| Control vs 24PD | 1 | 0.859 | 6.53 | 0.179 | **0.01** |
| Control vs LT25 | 2 | 0.047 | 0.28 | 0.017 | 1 |
| Control vs DS | 1 | 0.211 | 1.94 | 0.059 | 1 |
| Control vs LS | 1 | 0.125 | 1.41 | 0.042 | 1 |
| 24PD vs LT25 | 1 | 0.597 | 3.68 | 0.187 | **0.04** |
| 24PD vs DS | 1 | 0.266 | 1.21 | 0.074 | 1 |
| 24PD vs LS | 2 | 0.552 | 1.50 | 0.167 | 0.1 |
| LT25 vs DS | 1 | 0.130 | 1.09 | 0.060 | 1 |
| LT25 vs LS | 1 | 0.077 | 0.94 | 0.050 | 1 |
| DS vs LS | 1 | 0.148 | 1.14 | 0.063 | 1 |

***Beauveria ITS***
| Pairwise comparisons | Df | SumsOfSqs | F.Model | R2 | P-value |
| --- | --- | --- | --- | --- | --- |
| Control vs 24PD | 1 | 0.321 | 1.027 | 0.043 | 1 |
| Control vs LT25 | 2 | 0.284 | 0.446 | 0.036 | 1 |
| Control vs LS | 1 | 0.284 | 0.896 | 0.037 | 1 |
| Control vs DS | 1 | 0.370 | 1.195 | 0.047 | 1 |
| 24PD vs LT25 | 1 | 0.274 | 0.771 | 0.072 | 1 |
| 24PD vs LS | 1 | 0.286 | 0.714 | 0.082 | 1 |

|  |  |  |  |  |  |
| --- | --- | --- | --- | --- | --- |
| 24PD vs DS | 1 | 0.309 | 0.833 | 0.085 | 1 |
| LT25 vs LS | 1 | 0.285 | 0.782 | 0.073 | 1 |
| LT25 vs DS | 1 | 0.332 | 0.967 | 0.081 | 1 |
| LS vs DS | 1 | 0.290 | 0.759 | 0.078 | 1 |

